# Structural basis for inefficient G protein coupling by Frizzled receptors

**DOI:** 10.64898/2026.11.03.763784

**Authors:** Wessel A.C. Burger, Susovan Das, Hariprasad Venugopal, Ada S. Quinn, Billy J. Williams-Noonan, Lilian L. L. Wong, Tin T. Nguyen, Joshua R. Prendergast, Nicholas S. Kirk, Andrew P. Leis, Marija Dramicanin, Tracy Josephs, Matthew E. Call, Gunnar Schulte, Megan L. O’Mara, Kevin Huynh, Alisa Glukhova

## Abstract

Frizzleds (FZDs) are class F G protein-coupled receptors (GPCRs) that primarily signal through Dishevelled (DVL) to initiate Wnt signalling, with evidence for context-specific coupling to heterotrimeric G proteins. Every FZD–G protein structure to date has been determined without an agonist, implying that agonist binding is not required for stable complex formation *in vitro* and directly contradicting the difficulty of demonstrating G protein coupling in cells. All these structures contain unassigned density between the receptor and the heterotrimeric G protein. From structures generated by single-particle cryo-electron microscopy (cryo-EM) of FZD7–Gs and FZD5–Gq, we identify this density as the detergent lauryl maltose neopentyl glycol (LMNG), which bridges the receptor and G protein to stabilise a weak, shallow interface. Absence of LMNG — in biochemical preparations and molecular dynamics simulations — destabilises the complex, showing that constitutive FZD–G protein engagement is intrinsically weak. The weak G protein coupling is governed by a conserved intracellular interface that when mutated differentially influences DVL and G protein signalling. Our findings show that LMNG has masked the instability of purified FZD–G protein complexes and provide the structural basis of FZDs as inherently weak G protein activators that are predisposed to DVL signalling.

## Introduction

Frizzleds (FZDs) are a family of cell-surface receptors that mediate Wnt signalling, a pathway essential for embryonic development, tissue homeostasis, and disease^1,2^. FZDs are unusual members of the G protein-coupled receptor (GPCR) superfamily. They share the family’s classic seven-transmembrane (TM) architecture and are classified as class F GPCRs. Nevertheless, the main Wnt-induced and FZD-mediated signalling pathway, Wnt/β-catenin signalling, does not require heterotrimeric G proteins^3^. Instead, FZDs, together with co-receptors such as low-density lipoprotein receptor-related protein 5 and 6 (LRP5/6), bind Wnt glycoproteins and mediate Wnt/β-catenin signalling through activation of the intracellular scaffold protein Dishevelled (DVL)^4–6^.

Evidence for Wnt-induced G protein signalling at FZDs is long-standing but has been difficult to consolidate^7^. Early studies demonstrated that Wnt signalling mobilises intracellular calcium, a response disrupted by G protein inhibition^8–10^, and later work has described the potential role of heterotrimeric G proteins in FZD signalling^11–18^.

In contrast to this functional evidence, cryo-electron microscopy (cryo-EM) structures of FZD–G protein complexes provide direct demonstration that FZDs are capable of G protein coupling^19,20^. Remarkably, these structures resemble the architecture of classical GPCR–G protein complexes, including the outward displacement of TM helix 6 and the engagement of the G protein Cα5 helix with the receptor TM core. The structural similarity and the relative biochemical ease with which these complexes were obtained is somewhat surprising given the historical difficulty in demonstrating G protein coupling in cells. Additionally, these structures do not contain a bound agonist that is normally required to stabilise GPCR-G protein complexes according to the ternary complex model^21^.

It remains unclear whether the observed G protein coupling in these structures reflects an intrinsic, constitutive property of FZDs, or is instead a consequence of the biochemical preparations required for structural studies. Moreover, one striking feature observed in every FZD-G protein structure to date is unassigned density between the receptor and the bound G protein, positioned within the intracellular cavity of the FZD TM bundle and above the Cα5 helix of the Gα subunit^19,20,22^.

To address these questions, we determined the cryo-EM structures of FZD5–Gq and FZD7–Gs complexes at 2.7 Å and 2.6 Å respectively, improving on previous reconstructions at the FZD–G protein interface. We identified the density as the detergent lauryl maltose neopentyl glycol (LMNG) used during purification, and showed that it bridges the receptor and G protein. Here it stabilises the complex for structural studies, but in doing so obscures the intrinsic instability of FZD-G protein interactions. Moreover, we demonstrated that a conserved intracellular interface in FZDs underlies this weak G protein engagement and instead favours DVL coupling. Altogether, our results enhance our understanding of how FZDs couple to transducer proteins and shed light on why FZDs are unique within the GPCR family.

## Results

### Unique G protein engagement is observed at FZD7 and FZD5

As all FZD-G protein structures determined to date represent constitutively active FZDs, we asked how efficiently wild type (WT) FZD7 engages and activates Gs in the absence of agonist compared to the β2-adrenergic receptor (β_2_AR), a class A GPCR. Cells were semi-permeabilised with digitonin and we measured direct Gs heterotrimer coupling following treatment with apyrase, and G protein dissociation following the addition of GDPβS^23^. Compared to the β_2_AR, FZD7 displayed a substantially reduced capacity to couple to Gs (Fig. 1a) and trigger Gs dissociation (Fig. 1b-c), despite comparable receptor expression levels (Fig. 1d). This poor ability to couple and signal through G proteins is somewhat surprising given the stability of agonist-free FZD-G protein complexes used for structural studies^19,20^.

**Fig. 1.**
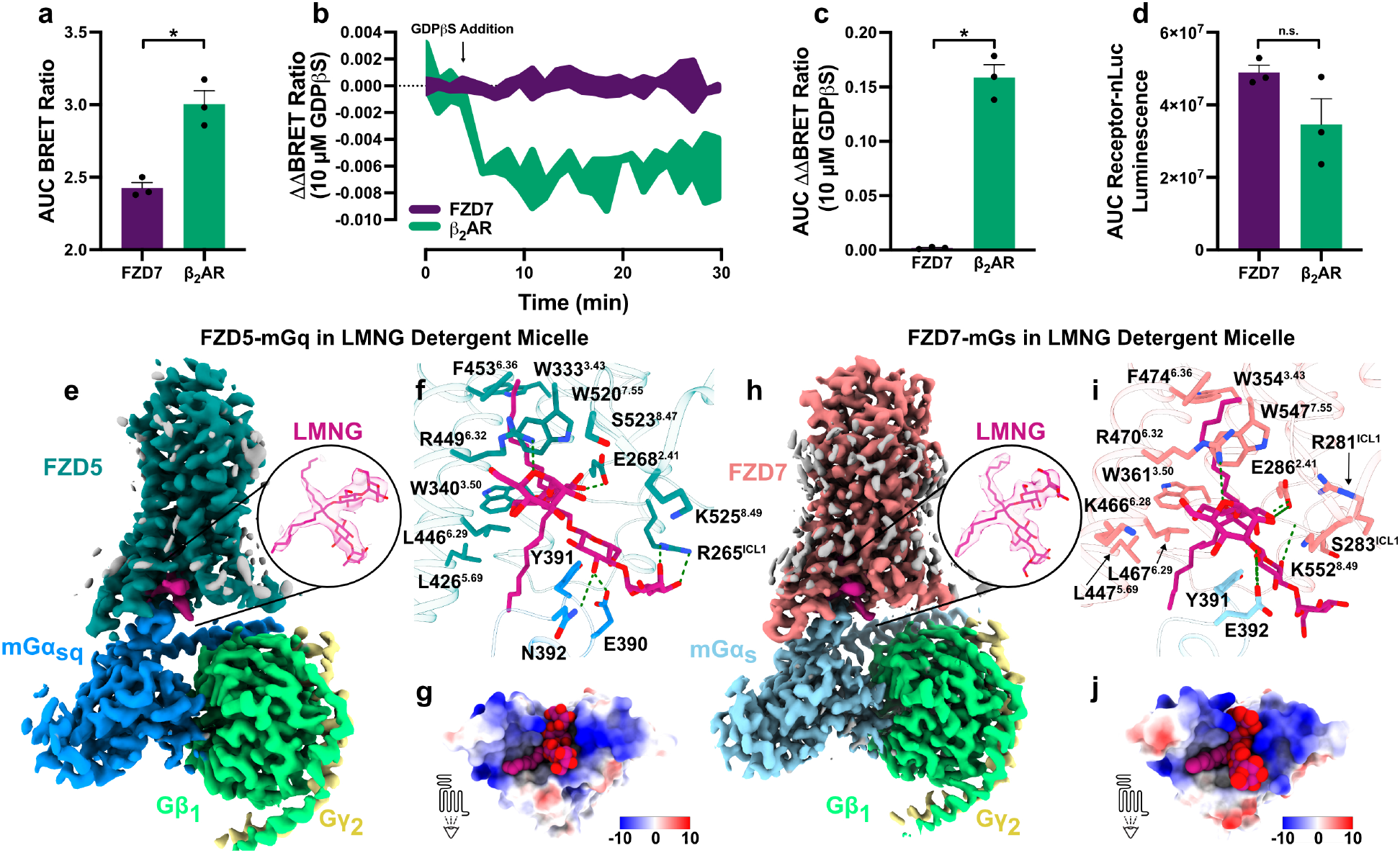
Weak FZD–G protein coupling is stabilised by a bridging detergent molecule in cryo-EM structures. **a** Gs coupling at WT FZD7 and β_2_AR, measured by BRET in digitonin-permeabilised and apyrase treated ΔFZD1-10 HEK293T cells transfected with receptor-nanoluciferase, WT Gα_sL_, Gβ_1_, Gγ_2_-Venus. AUC of the BRET ratio over 30 min after NanoGlo substrate addition. **b** Representative kinetic ΔΔBRET trace showing Gs dissociation following addition of 10 μM guanosine 5′-[β-thio]diphosphate (GDPβS). **c** AUC of the ΔΔBRET traces shown in b. **d** AUC of receptor-nLuc luminescence, showing comparable receptor expression between FZD7 and β_2_AR. Bars in a,c-d represent mean ± s.e.m. of n=3 independent biological experiments, each performed in duplicate. *, significantly different, *P <* 0.05, Welch’s t test. **e** Cryo-EM density map of the FZD5–mGq complex in an LMNG detergent micelle; inset shows LMNG density. **f** LMNG in the FZD5 intracellular cavity, with LMNG and contacting residues shown as sticks, hydrogen bonds shown as green dashes. **g** Electrostatic surface of the FZD5 intracellular cavity viewed from the membrane, LMNG shown as spheres. **h-j** As in e-g for the FZD7-mGs complex in LMNG micelle. Composite map and LMNG density contoured at 8.27, σ = 6.64. Source data are provided as a Source Data file.

In all four FZD–G protein complexes determined to date, prominent and unassigned electron density caps the Cα5 helix of the Gα subunit^19,20,22^ (Supplementary Fig. 1a-d). To identify this density and to clarify the mechanism of G protein-coupling at FZDs, we determined the high-resolution structures of FZD5 and FZD7 coupled to heterotrimeric G proteins. Both receptors constitutively activate multiple G protein subtypes in the absence of Wnt agonists (Supplementary Fig. 2). FZD5-Gq signalling has been implicated in the survival of pancreatic cancer cells^11^, and FZD7-Gs signalling in skeletal muscle hypertrophy^24^. We therefore focused on Gq and Gs at FZD5 and FZD7, respectively. We used cryo-EM to determine the structures of FZD5-mGq and FZD7-mGs in LMNG detergent to global resolutions of 2.7 Å and 2.6 Å, respectively (Supplementary Fig. 3,4).

Overall, our structures resemble the previously reported ligand-free FZD-G protein complexes^19,20,22^. The extracellular cysteine-rich domain (CRD) of FZD is flexible and is unresolved at the reported resolution. The extracellular loops form a tight bundle above the TM domain, which itself displays the hallmarks of active-state GPCRs. The TM6 kinks at P^6.43^ (superscript refers to the Ballesteros and Weinstein scheme for GPCR residues^25^) to open an intracellular cavity for G protein binding. However, the G protein engages FZD far more superficially compared to other GPCR–G protein complexes, including that of the closest FZD relative Smoothened (SMO) (Supplementary Fig. 1e). The Cα5 helix of Gα docks against the intracellular cavity rather than inserting into it, with its C-terminal residue positioned at the cavity entrance. Because of this shallow G protein engagement, the receptor–Gα interface is only 546 Å^2^ for FZD7-mGs and 496 Å^2^ for FZD5-mGq, compared to 958 Å^2^ SMO–Gi or 1260 Å^2^ in β_2_AR-Gs structures (Supplementary Fig. 1f,g). This shallow, low-surface area engagement is found at all available FZD-G complexes^19,20^. As these consist of different FZD and G protein subtypes, it indicates that this is a general feature of FZD–G protein coupling rather than a specific property of any single FZD receptor or G protein.

In both our FZD7-Gs and FZD5-Gq structures, the intracelullar cavity is occupied by a large density that cannot be assigned to either receptor or G protein (Fig. 1e, h) and is similar to the density observed in all FZD-G protein structures determined to date (Supplementary Fig. 1a-d). The density resembles two elongated, lipid-tail-like moieties linked to what appear to be two headgroups composed of relatively planar ring structures. One tail projects vertically into the TM bundle (TMs 2, 3, 5-7) while the other extends toward the cytoplasmic face of TMs 3, 5, and 6. Both headgroups protrude toward the solvent - one positioned at the TM7-helix 8 kink and the other stacked in between the top of the Cα5 helix and receptor intracellular loop (ICL) 1 - and are surrounded by positively charged residues (Fig. 1g, j).

The presence of two distinct tails and two headgroups is inconsistent with an endogenous lipid. Since all FZD-G protein complexes determined by cryo-EM to date were purified in LMNG, we assigned this density as LMNG, which provided an excellent fit to the experimental map in both FZD5 and FZD7 structures (Fig. 1e, h). The first lauryl tail is wedged into the TM core occupying a hydrophobic groove lined by W^3.43^, F^6.36^ and W^7.55^ (Fig. 1f, i). The second LMNG lauryl tail, wedged towards the bottom of TM3 and the Cα5 helix of the G protein, engages with L^5.69^ and L^6.29^ of both FZD5 and FZD7 and with the Cα5 helix (H387, L388, Y391 and L393 of Gα_s_ and N387, L388, Y391 and L393 of Gα_q_) through Van der Waals interactions. At both FZD5 and FZD7, the maltose headgroups of LMNG form hydrogen bonds with E^2.41^ and R^6.32^. At FZD5, LMNG displays additional hydrogen bonds with R265 of ICL1 and with residue E390 of Gα_q_ (Fig. 1f). At FZD7, LMNG instead forms an additional hydrogen bond with the Gα residue E392 and with the backbone of Y284^2.39^ (Fig. 1i). Through these interactions LMNG increases the FZD-G protein contact area from 546 Å^2^ to 686 Å^2^ for FZD7 and 496 Å^2^ to 609 Å^2^ for FZD5. Given the likely presence of LMNG in all reported active-state FZD structures, we considered whether LMNG: 1) directly perturbs G protein binding, leading to shallow engagement; 2) stabilises an otherwise transient complex; 3) passively fills space at a weak protein-protein interface; or 4) substitutes an endogenous lipid.

### LMNG stabilises the complex for cryo-EM

To address whether LMNG perturbs G protein binding at FZDs, we determined the structure of FZD7-mGs reconstituted into lipid nanodiscs (Fig. 2a, Supplementary Fig. 5). FZD7-mGs was solubilised and purified using n-dodecyl-β-D-maltoside (DDM) which we anticipated has a lower affinity for the intracellular cavity than LMNG because of its single lipid tail and a single headgroup. Following reconstitution into POPC/POPG MSP1D1 nanodiscs, the structure was solved to a global resolution of 2.8 Å. The nanodisc-reconstituted FZD7-mGs structure closely resembled the LMNG-solubilised structure (RMSD 0.9 Å) (Fig. 2c). Strikingly, a similar density was again observed at the FZD-G protein interface, matching the position and shape of the density seen in the LMNG structure (Fig. 2b). Mass spectrometry of two additional samples generated with the same reconstitution protocols detected 100 and 150 pmol of residual per µg of protein — a molar ratio of 22:1 and 33:1, respectively, relative to the FZD7–mGs complex — most likely carried over from unrelated purifications performed in the laboratory (Supplementary Fig. 5). Strikingly, these amounts of LMNG were sufficient to fully occupy the intracellular cavity, suggesting high-affinity binding to FZD7.

**Fig. 2.**
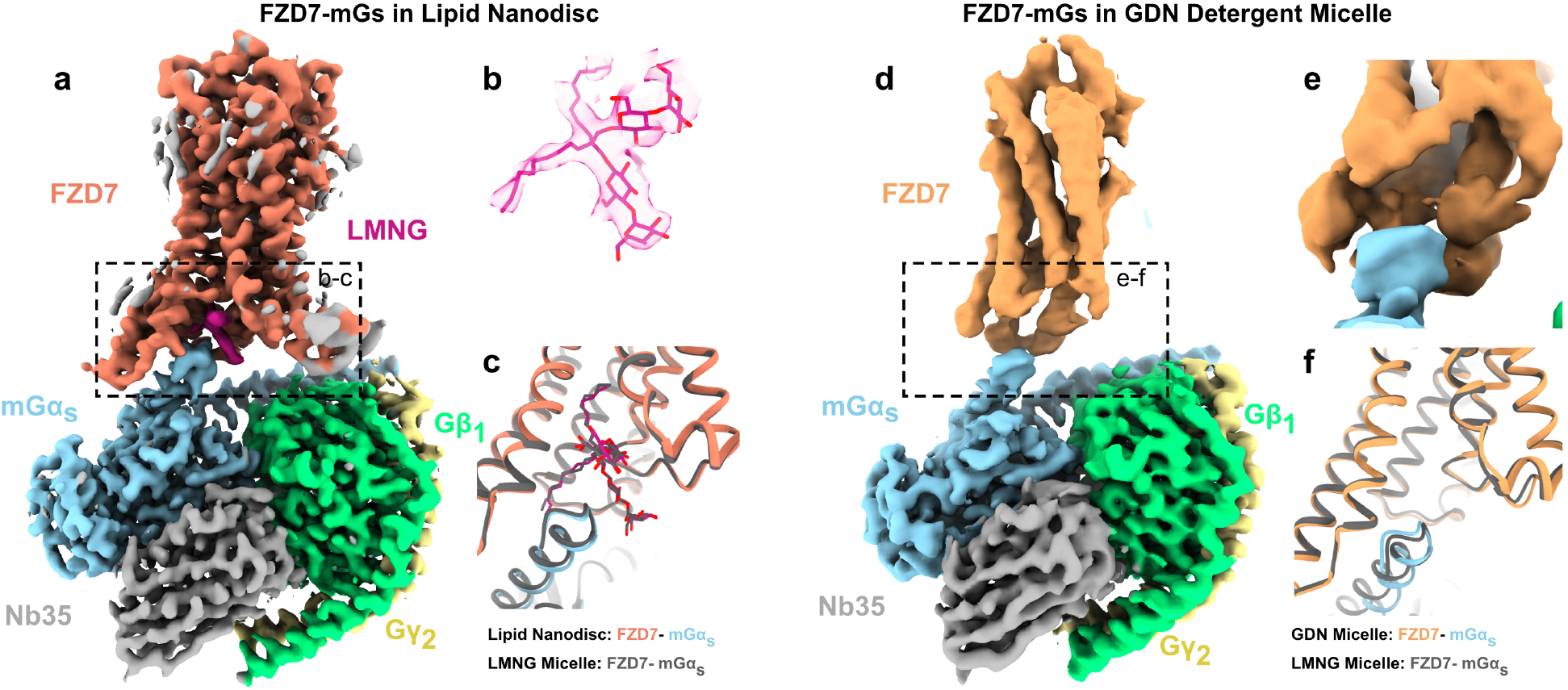
FZD7-mGs adopts the same architecture in lipid nanodisc and in GDN micelle. **a** Cryo-EM density map of the FZD7-mGs complex reconstituted in MSP1D1 lipid nanodisc. Composite map and LMNG density contoured at 8.6, σ = 7.74. **b** Density for the modelled LMNG molecule shown as sticks. **c** Comparison of the nanodisc structure with the LMNG micelle structure from Fig. 1h (grey), shown as ribbons. **d** Cryo-EM density map of the FZD7-mGs complex in glyco-diosgenin (GDN) detergent micelle, contoured at 0.0299, σ = 8.00. **e** Close-up of the FZD7-mGα_s_ interface in the GDN map, contoured at 0.0247, σ = 6.61; no ordered lipid or detergent density is present at the site occupied by LMNG in the lipid nanodisc complex. **f** Superposition of the GDN micelle structure with the LMNG micelle structure from Fig. 1h.

To eliminate LMNG entirely, we purified FZD7–mGs in an independent laboratory with no prior use of LMNG and confirmed all reagents were LMNG-free by mass spectrometry. The complex was extracted in DDM and exchanged into glyco-diosgenin (GDN) during purification. Complex formation was verified by size-exclusion chromatography and Coomassie staining prior to grid preparation (Supplementary Fig. 6). Cryo-EM imaging, however, revealed that the majority of complexes dissociated during vitrification. 20,585 micrographs gave 26,563,686 picked particles whilst only 78,395 particles (0.3%) contributed to the final 3.1 Å reconstruction, compared with 10% for the LMNG dataset. The map quality supported modelling of the G protein side chains, but only rigid-body fitting of the FZD7 TM core (Fig. 2d, Supplementary Fig. 6). In this structure, the G protein retained the same shallow engagement with FZD7, seen in our micelle and nanodisc structures and all FZD-G complexes reported to date (Fig. 2f). Critically, the intracellular cavity remained empty, even at increased contour levels (Fig. 2e).

These results indicate two things. First, the shallow engagement of G proteins by FZDs is not an LMNG artefact: it is present in LMNG micelles, lipid nanodiscs, and an LMNG-free GDN preparation. Second, the binding of LMNG during complex purification has a pronounced stabilising effect.

### MD simulations reveal that LMNG stabilises a weak FZD – G protein interaction

To examine how LMNG affects the dynamics of the FZD7–Gs complex, we performed molecular dynamics (MD) simulations of the LMNG-bound FZD7–mGs structure embedded in a POPC/cholesterol bilayer, with and without the modelled LMNG molecule. Each condition was simulated as 3 independent replicas of 500 ns from identical starting coordinates and randomised initial velocities. All trajectories remained stable throughout (Supplementary Fig. 7a) and with LMNG bound, the complex remained in an active conformation. With LMNG removed, however, the trajectories revealed reduced FZD7-mGs complex stability, consistent with early steps of complex dissociation^26^. Specifically, the G protein Cα5 helix was destabilised (Supplementary Fig. 7b) and began to unfold and rotate inward, bringing Y391^Gαs^ and R38^Gαs^ into proximity to form a cation–π interaction characteristic of the GDP-bound Gα_s_^27–29^ (Fig. 3a). Concurrently, the receptor exhibited conformational changes associated with receptor inactivation, including closure of the TM domain as measured by the distance between the L467^6.29^ and G551^8.48^ Cα atoms (Fig. 3b). These changes were not observed in the LMNG-bound simulations.

**Fig. 3.**
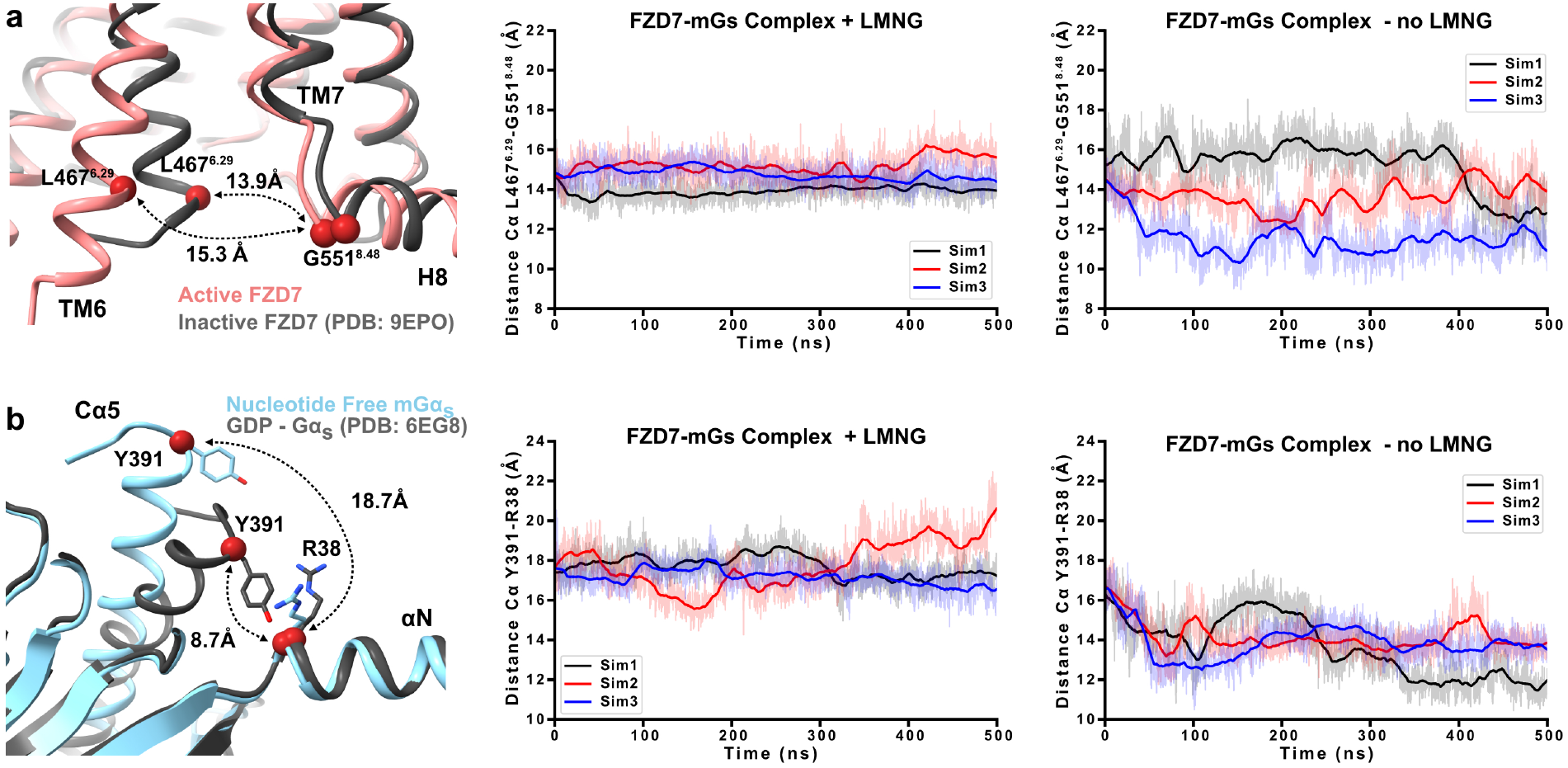
LMNG stabilises a weak FZD – G protein interaction in molecular dynamics simulations. **a** Left, comparison of the active FZD7 structure determined here with inactive FZD7 (PDB 9EPO), showing the Cα-Cα distance between L467^6.29^ and G551^8.48^ : 15.3 Å in the active state and 13.9 Å in the inactive state. Middle and right, the L467^6.29^-G551^8.48^ Cα-Cα distance over 500 ns of molecular dynamics (MD) simulation of the FZD7-mGs complex with LMNG bound (middle) and after removal of LMNG (right). Three independent trajectories per condition are shown in different colours. **b** Left, superposition of nucleotide-free mGα_s_ from the FZD7-mGs structure with GDP-bound Gα_s_ (PDB 6EG8), showing the Cα-Cα distance between Y391 (Cα5 helix) and R38 (αN helix): 18.7 Å in the nucleotide-free state and 8.7 Å in the GDP-bound state. Middle and right, the Y391-R38 Cα-Cα distance over 500 ns of MD simulation with LMNG (middle) and without LMNG (right). Three independent trajectories per condition are shown in different colours.

Altogether, the simulations indicate that the FZD7–Gs interaction is not stable in the absence of LMNG as the complex transitions away from the nucleotide-free, receptor-bound state. Similarly, MD simulations of LMNG-bound FZD7 in the absence of G protein revealed increased LMNG mobility within the binding site compared to the G protein-bound conditions, indicating that the G protein reciprocally stabilises LMNG binding to the receptor (Supplementary Fig. 7c). These reciprocal effects point to an allosteric interaction between LMNG, FZD, and G protein, whereby LMNG acts as a molecular glue to stabilise the FZD7–G protein complex.

### Conserved residues underlie poor FZD-G protein activation

Our structural and MD data reveal that FZDs engage G proteins through a shallow interface, in stark contrast to SMO, where the Cα5 helix fully occupies the intracellular cavity (Supplementary Fig. 1e,g). We hypothesised that differences in the amino acid residues lining the intracellular binding site are responsible for the contrast between robust, agonist-induced G protein coupling at SMO^30,31^ and the poor G protein binding observed for FZDs (in the absence of LMNG), as well as for the FZD preference for DVL^32^. The residues at this site are highly conserved amongst all ten FZDs (Fig. 4a). We identified four residues in SMO (A^2.41^, F^3.54^, E^6.28^, T^6.29^) that form direct contacts with the G protein but differ in FZDs (E^2.41^, A^3.54^, K^6.28^, L^6.29^). These residues engage with LMNG in the intracellular cavity, but form no direct contact with the G protein (Fig. 4a-c). We hypothesised that swapping these residues to their SMO counterparts would enhance the G protein coupling at FZDs. To test this, we generated E286^2.41^A, A365^3.54^F, K466^6.28^E, L467^6.29^T and a “Quad” (all four mutations) FZD7 mutants.

**Fig. 4.**
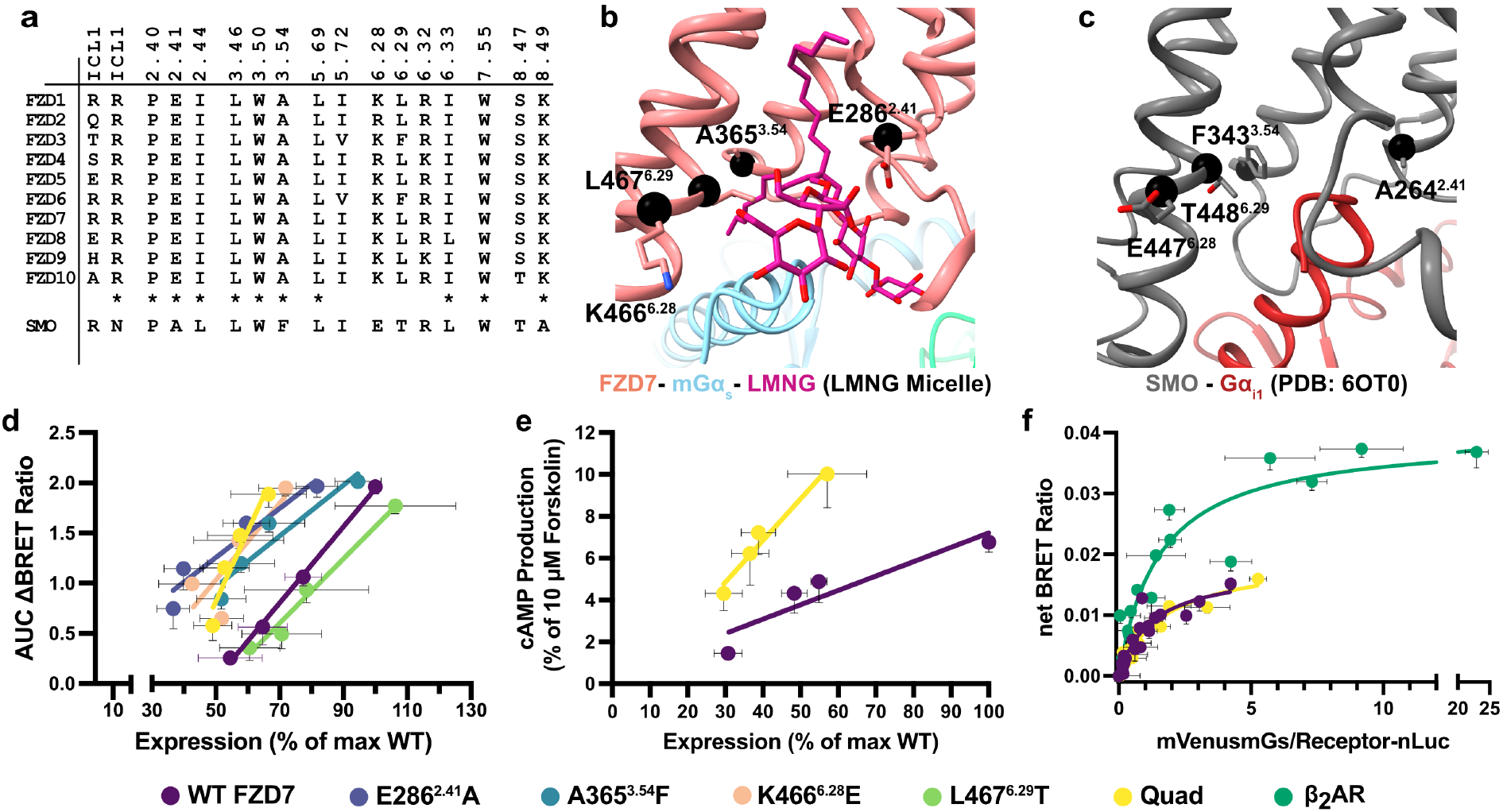
Mutation of the highly conserved intracellular cavity increases constitutive G protein activation and cAMP production. **a** Sequence alignment of the cavity-lining residues across human FZD1-10 subtypes and Smoothened (SMO). Asterisks mark positions fully conserved across the FZD family. **b** LMNG (sticks) bound in the FZD7–mGs structure; black spheres mark the Cα atoms of the four mutated positions. **c** Equivalent region of the SMO-Gi1 structure (PDB 6OT0), with the Cα atoms of the corresponding SMO residues shown. **d** Constitutive Gs activation (AUC of ΔBRET ratio) versus receptor expression (% of max WT FZD7) in ΔFZD1-10 HEK293T cells transfected with increasing WT or mutant FZD7 DNA. Data points represent mean ± s.e.m., of n = 4 independent experiments (ΔBRET ratio performed in duplicate; expression performed in triplicate); lines represent linear regression fit. **e** Constitutive cAMP production (% forskolin response) versus receptor expression (% of max WT FZD7) for WT FZD7 and Quad in ΔFZD1-10 HEK293T cells transfected with increasing WT or Quad DNA. Data points represent mean ± s.e.m., n = 5 experiments performed in triplicate; lines represent linear regression fits. **f** BRET saturation assay of mGα_s_ recruitment. ΔFZD1-10 HEK293T cells were transfected with constant receptor-nLuc (donor) and increasing mVenus-mGα_s_ (acceptor). Net BRET ratio is plotted against mVenus fluorescence/nLuc luminescence ratio. Data points represent mean ± s.e.m. (both axes), pooled from n = 4 (WT) and n = 3 (Quad, β_2_AR) experiments performed in duplicate. Curves represent the fit of a one-site specific binding model. BRET_50_ and BRET_max_ values listed in Supplementary Table 3. Source data are provided as a Source Data file.

We used the TRUPATH^33^ assay to compare constitutive Gs activation between different FZD7 constructs. We transfected increasing amounts of receptor and measured Gs heterotrimer activation and quantified total expression (intracellular + cell-surface) by flow cytometry to control for differences in receptor expression (Supplementary Fig. 8a). For all constructs, G protein activation increased with receptor expression, indicative of constitutive activity (Fig. 4d). Porcupine inhibition, which blocks endogenous Wnt secretion, did not affect Gs activation, ruling out receptor activation from endogenous Wnts^34,35^ (Supplementary Fig. 8b). While L467^6.29^T displayed similar activity to WT FZD7, the remaining single mutants and the Quad mutant all showed enhanced activity, displaying higher activation at comparable receptor expression levels. This effect was most striking in the Quad mutant, where small increases in expression led to large increases in Gs activation (Fig. 4d). Because TRUPATH is performed with overexpressed G proteins and highly optimised bioluminescence resonance energy transfer (BRET) sensors, we asked whether the enhanced constitutive activity of the Quad mutant persists with endogenous G proteins. Use of the CAMYEL biosensor to detect changes in intracellular cAMP levels showed enhanced constitutive activity at the Quad mutant, consistent with the TRUPATH results (Fig. 4e).

To determine whether the enhanced constitutive activity is due to greater G protein coupling at the Quad mutant, we performed saturation net BRET assays measuring mVenus-mini-Gα_s_ (mGα_s_) binding to FZD7-nLuc. Comparable G protein binding was observed at both WT FZD7 and the Quad mutant, with similar BRET_50_ apparent affinities (WT: 1.25 ± 0.27, Quad: 1.69 ± 0.26) and BRET_max_ values (WT: 0.018 ± 0.0018, Quad: 0.020 ± 0.0015) (Fig. 4f). Of note, the β_2_AR displayed higher BRET_max_ values (0.040 ± 0.0023), further highlighting the intrinsically limited G protein coupling capacity of FZD7. mGα_s_ lacks a functional nucleotide-binding pocket and is therefore incapable of nucleotide exchange or receptor dissociation, limiting its ability to resolve potential differences in steady-state G protein turnover. We therefore turned to mVenus-tagged WT Gα_s_, which revealed similar BRET_50_ apparent affinities and BRET_max_ values at both WT FZD7 and the Quad mutant (Supplementary Fig. 8d, Supplementary Table 3).

Our G protein coupling and signalling data suggest that shallow Cα5 insertion renders FZDs inherently inefficient at G protein activation. This deficiency can be overcome by introducing SMO-like residues at the intracellular interface. To understand the structural basis for this functional rescue, we determined the structure of the Quad mutant by cryo-EM.

### Quad mutant structure reveals an empty intracellular cavity and a remodelled G protein interface

We determined the structure of the FZD7 E286^2.41^A/A365^3.54^F/K466^6.28^E/L467^6.29^T Quad mutant in complex with mGs heterotrimer in LMNG detergent micelle at a global resolution of 3.0 Å (Fig. 5a, Supplementary Fig. 9). The Quad structure closely resembles that of WT FZD7 in both LMNG and nanodiscs (RMSD of 1.3 Å and 1.8 Å, respectively). Notably, despite purification in LMNG, no electron density was observed within the intracellular cavity (Fig. 5b). As with WT FZD7, the G protein remained shallowly bound (Fig. 5c). Clear side-chain density was observed for the A365^3.54^F and L467^6.29^T substitutions, explaining how LMNG binding is disrupted. The A365^3.54^F side chain occupies the position of the lauryl tail within the TM core, while L467^6.29^T introduces a polar hydroxyl group (Fig. 5d). Additionally, four non-mutated residues adopt different rotamers in the Quad mutant structure compared to the WT (Fig. 5d). W361^3.50^, F474^6.36^, W547^7.55^ and H371^4.35^ rotate into positions that clash with either the second lauryl chain or the headgroup of LMNG. Collectively, the changes in the Quad mutant induce a similar conformation at the receptor intracellular side as seen at the agonist-bound SMO receptor (Fig. 5e) as well as a larger contact area between the receptor and G protein (647 Å^2^) compared to the WT FZD7 (546 Å^2^, omitting the LMNG from the structure, Supplementary Fig. 1f).

**Fig. 5.**
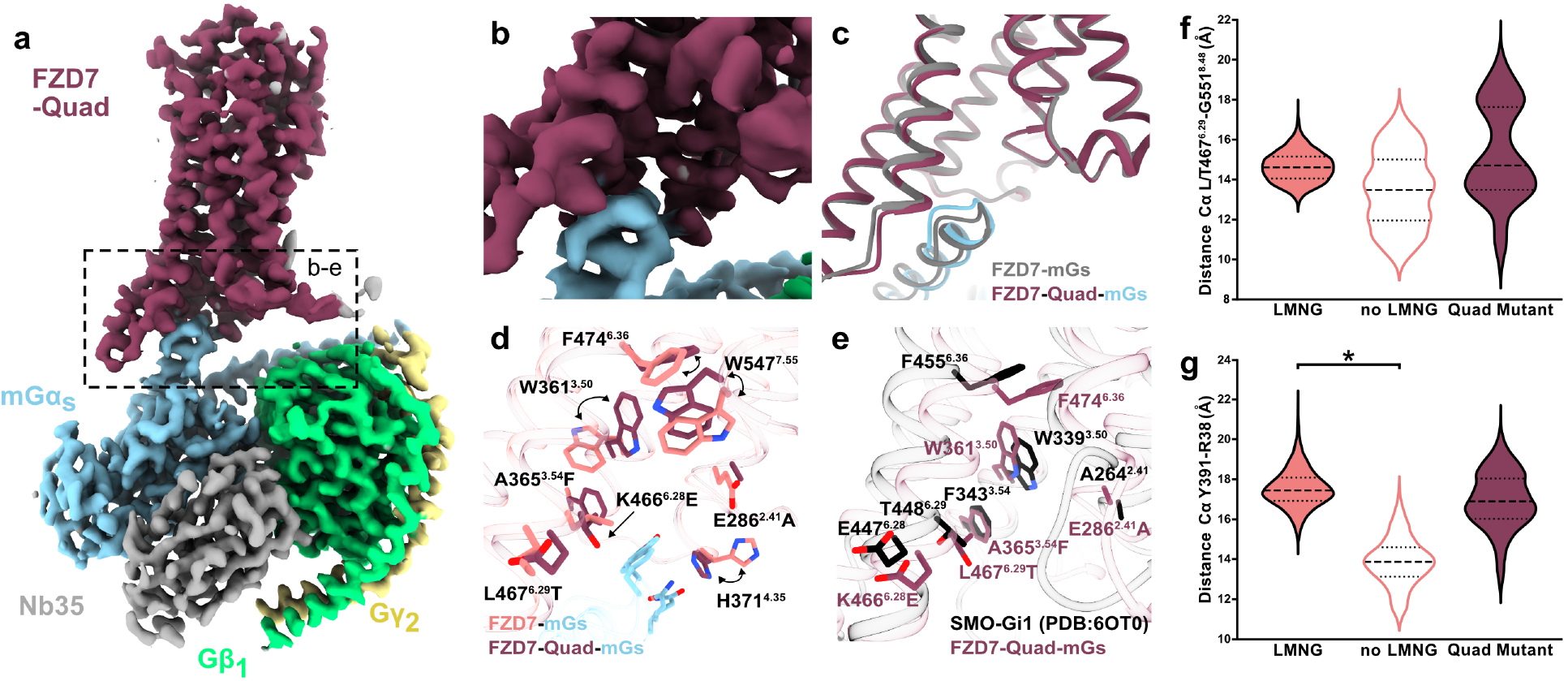
Cryo-EM structure of Quad mutant reveals a rearranged G protein interface that stabilises G protein interaction. **a** Cryo-EM density composite map of the FZD7-Quad-mGs complex in LMNG micelle. Map contoured at 7.45, σ = 7.03. **b** Close-up of the FZD7-Quad/mGα_s_ interface; no density is present at the site occupied by LMNG in the WT complex. **c** Superposition of FZD7-Quad-mGs with WT FZD7-mGs shown as cartoons. **d** Side-chain rearrangements of key residues in FZD7-Quad-mGs shown as sticks compared to WT FZD7-mGs. Arrows indicate the repositioning of W361^3.50^, F474^6.36^, W547^7.55^ and H371^4.35^ that accompanies the E286^2.41^A, A365^3.54^F, K466^6.28^E and L467^6.29^T substitutions. **e** Superposition of FZD7-Quad-mGs with SMO-Gi1 (PDB 6OT0), showing the equivalent SMO residues A264^2.41^, F343^3.54^, E447^6.28^, T448^6.29^, W339^3.50^ and F455^6.36^. **f-g** Distributions of the Cα-Cα distance between residue 467^6.29^ (L in WT, T in Quad) and G551^8.48^ **(f)** and between Y391 (Cα5 of Gα_s_) and R38 (αN of Gα_s_) **(g)**, sampled over 500 ns of MD simulation of the FZD7-mGs complex in the presence and absence of LMNG and for the Quad mutant. Data for LMNG and no LMNG are replotted from Fig. 3a,b. Each violin pools three independent trajectories per condition (n = 3), giving 15,000 frames per distribution and shows a kernel density estimate of the pooled values. Dashed lines mark the median and dotted lines the first and third quartiles. *, significantly different, *P <* 0.05, Welch’s t test on the per-trajectory means, unmarked comparisons were not significant (*P* > 0.05).

We next used MD simulations to assess dynamics of the complex. The TM domain of the Quad mutant exhibited greater conformational variability, including increased opening, relative to both the LMNG-bound and LMNG-free WT complexes (Fig. 5f), which may facilitate G protein coupling and insertion. In contrast, the Quad mutant preserved the outward rotation of the Gα_s_ Cα5 helix, with the Y391^Gαs^ and R38^Gαs^ distance remaining close to that of the LMNG-bound WT complex rather than collapsing as observed for the LMNG-free WT complex (Fig. 5g). Together, these results suggest that the Quad mutations and associated rotameric changes compensate for the loss of LMNG, recapitulating the reduced dynamics of the LMNG-bound WT complex.

The structural and dynamic properties of the Quad mutant provide a basis for its enhanced constitutive activity: the mutations mimic the conformations observed at a GPCR fully engaged to a G protein (like SMO) leading to enhanced complex stability in MD simulations and improved G protein signalling (Fig. 4). In the WT FZD7 complex, this is achieved through LMNG binding during protein purification. However, in the absence of LMNG, such as in a cellular context, FZDs remain inherently poor G protein couplers.

### A conserved interface underlies the mechanism of transducer coupling at FZDs

We next examined whether the conserved intracellular interface of FZDs is responsible for their preferential coupling to DVL. TOPFlash assays that measure β-catenin-dependent transcription were used to capture DVL activation. FZD7 exhibits limited DVL activity in the absence of Wnt stimulation (Supplementary Fig. 8c); therefore, we stimulated cells with Wnt3a. All mutations impaired DVL-dependent Wnt/β-catenin signalling, with complete loss of signalling observed at the L467^6.29^T, A365^3.54^F and Quad mutants (Fig. 6a). Saturation net BRET assays measuring mVenus-DVL2 recruitment to FZD7-nLuc revealed that the Quad mutations abolish DVL2 coupling, with non-specific binding levels indistinguishable from those of the β_2_AR negative control that does not engage DVL (Fig. 6b).

**Fig. 6.**
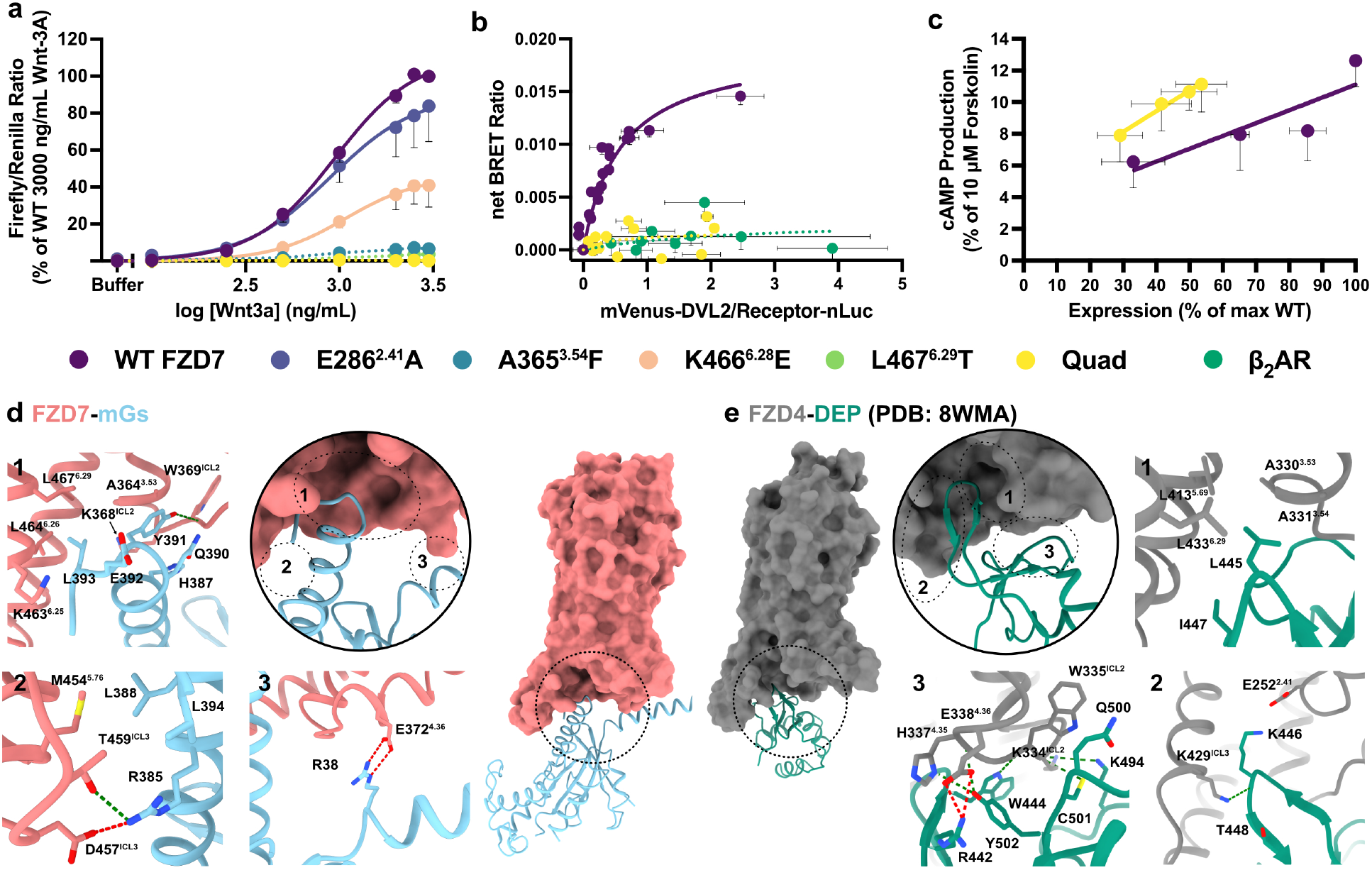
Quad mutations uncouple FZD7 from Wnt3a-induced β-catenin signalling and DVL2 recruitment while retaining Gs coupling. **a** Wnt3a concentration–response curves in ΔFZD1-10 HEK293T cells expressing WT FZD7 or the indicated mutants. Data points represent mean ± s.e.m. of n = 4 for WT and E286^2.41^A, n = 3 for A365^3.54^F, K466^6.28^E, L467^6.29^T and Quad experiments performed in duplicate and normalised to response of 3,000 ng/mL Wnt3a observed at FZD7 WT. Curves represent four-parameter logistic fits. Dotted lines indicate mutants for which no fit could be determined. **b** BRET saturation assay of DVL2 recruitment. ΔFZD1-10 HEK293T cells were transfected with constant receptor-nLuc (donor) and increasing mVenus-DVL2 (acceptor). Net BRET ratio is plotted against mVenus fluorescence/nLuc luminescence ratio. Data points represent mean ± s.e.m. (both axes), pooled from n = 4 (WT) and n = 3 (Quad, β_2_AR) experiments performed in duplicate. Curves represent fit of a one-site specific binding model. Dotted lines indicate where no fit could be determined. Parameters obtained in a,b are listed in Supplementary Table 3. **c** Constitutive cAMP production (% forskolin response) versus receptor expression (% of max WT) for WT FZD7 and Quad in ΔDVL1-3 HEK293T cells transfected with CAMYEL biosensor and increasing WT or Quad DNA. Data points represent mean ± s.e.m. of n = 5 experiments performed in triplicate; lines represent linear regression fits. **d** FZD7-mGs interface with the boxed regions 1-3 shown as close-ups; hydrogen bonds are shown as green dashes, ionic bonds as red dashes. **e** Equivalent interface of the FZD4-DEP domain complex (PDB 8WMA), with matching close-ups 1-3. Source data are provided as a Source Data file.

Loss of DVL coupling at the Quad mutant raised the possibility that the enhanced G protein signalling (Fig. 4d,e) reflects, at least in part, reduced competition between G protein and DVL for a shared FZD7 binding interface. We tested this using ΔDVL1-3 HEK293T cells, reasoning that if competition alone could explain the increase, the Quad mutant should lose its enhanced constitutive activity in a DVL-null background. Instead, the Quad mutant retained higher constitutive activity in cAMP accumulation assays (Fig. 6c). Similarly, saturation net BRET assays with mVenus-WT Gα_s_ in ΔDVL1-3 HEK293T cells revealed similar apparent affinities (Supplementary Fig. 8e, Supplementary Table 3) indicating that the absence of DVL does not lead to enhanced G protein binding. These findings in ΔDVL1-3 HEK293T cells show that the G protein–FZD7 interaction is not influenced by DVL. Taken together, our data suggest that the conserved intracellular interface binds G proteins weakly whilst preferring DVL binding, indicating that this interface is a molecular determinant of FZD-transducer selectivity.

### Search for an endogenous lipid

To determine whether the intracellular cavity might represent an endogenous lipid binding site, we performed untargeted lipidomics on the WT FZD7-mGs complex purified in either LMNG or GDN, and the FZD7 Quad-mGs complex. We reasoned that an endogenous lipid occupying the pocket should be selectively enriched in the GDN-purified sample. If such lipids are present at substoichiometric quantities or are compositionally diverse, they would average out during cryo-EM data processing — appearing as an empty pocket — but should nonetheless be detectable by mass spectrometry. Conversely, the saturating presence of LMNG in the WT FZD7 complex, or the mutations that impair LMNG/lipid binding in the FZD7 Quad complex should serve as negative controls. Lipidomics revealed selective enrichment of lysophosphatidic acid (LPA) and a number of phosphatidylinositol monophosphate (PIP1) and phosphatidylinositol bisphosphate (PIP2) species in the WT FZD7 in GDN sample relative to both WT FZD7 in LMNG and FZD7 Quad mutant in LMNG samples (Supplementary Fig. 10).

To assess whether these lipids influence FZD7-mediated G protein activation, we performed constitutive activity assays using TRUPATH on both WT FZD7 and the Quad mutant. Because lipidomics identified specific acyl chain compositions for PIP1 and PIP2 that were not commercially available, we tested the closest available structural analogues, PIP1/PIP2 18:1 (equivalent to 36:2 for both chains), using PIP carriers for their in-cell delivery. None of the lipids tested altered constitutive FZD7 mediated G protein activation (Supplementary Fig. 10). This may indicate that the lipid composition differences observed in the lipidomics experiments arose from variation during protein extraction and purification. Alternatively, FZD7 might specifically associate with a narrow range of acyl chain phospholipids identified by lipidomics but unattainable for our validation experiments.

## Discussion

Our structures show that G proteins couple to FZDs in a shallow manner, leaving a large, unoccupied cavity within the receptor TM core. During routine preparations of FZD-G protein complexes for cryo-EM, this cavity is occupied with LMNG that artificially stabilises the complex. The presence of LMNG at the FZD–G protein interface is likely a feature of every FZD–G protein complex determined by cryo-EM to date^19,20^. LMNG likely binds FZD with high affinity, as it exhibited persistent occupancy following reconstitution into lipid nanodiscs, where trace carry-over detergent was sufficient to maintain full occupancy. Collectively, these observations mean that all previously determined active-state FZD structures should be reinterpreted with the understanding that LMNG acts as a molecular glue that bridges and stabilises an otherwise weak protein–protein interface. Our MD simulations support this as removal of LMNG from the FZD7–mGs complex initiates transitions towards the inactive conformations for both the G protein and the receptor. The shallow G protein engagement likely limits FZDs’ ability to activate heterotrimeric G proteins and may explain the complicated data for G protein coupling and signalling at FZDs in cells. These findings fundamentally challenge our interpretation of the growing structural record on class F signalling, and bring the structures into agreement with the long-standing difficulty of demonstrating FZD–G protein coupling and signalling in cells.

Concurrently, this weak G-protein engagement may also explain FZDs’ higher preference to signal through DVL^32^. Like the Cα5 helix of G proteins, the DVL DEP domain inserts shallowly into the FZD intracellular cavity. Despite this, the intracelullar cavity remains empty with no additional density present in the FZD4-DEP structure (despite use of LMNG during purification)^36^. The DEP domain and G proteins have a comparable surface area (648 Å^2^ for FZD4-DEP compared to 546 Å^2^ and 496 Å^2^ for FZD7-mGs and FZD5-mGq, respectively, Supplementary Fig. 1f) indicating that DVL and G proteins compete for a shared but non-identical binding surface. Both the finger loop and Cα5 helix of the DEP domain and G protein, respectively, display hydrophobic and hydrogen bonding interactions with the intracellular cavity and ICL3 of FZDs (Fig. 6d,e, Panels 1-2). However, the G protein’s αN helix displays a single hydrogen bond interaction with TM4 of FZD7 (R38-E372^4.36^, Fig. 6d, Panel 3), whereas the DEP domain displays multiple hydrogen and ionic bond interactions with TM4 and ICL2 of FZD4 (Fig. 6e, Panel 3). Furthermore, unlike G proteins, DVL is reported to engage a second interaction site between its PDZ domain and the KTXXXW motif in Helix 8 of FZDs^37,38^, potentially creating a bivalent FZD-DVL interface. The local concentration of DVL at the membrane may be further increased by FZD clustering^39^, DIX-domain-dependent polymerisation^40^ and by scaffolding within larger assemblies, including DVL–Axin interactions^41^. This multivalent engagement of DVL, combined with its increased local concentration at the FZD signalling complex, might represent a mechanism by which FZDs prioritise DVL coupling and signalling. Whether this simultaneously permits low-level G protein coupling remains to be determined. It is also plausible that certain cellular contexts, particularly those characterised by high G protein expression and low DVL concentrations, provide sufficient binding opportunities for G proteins to overcome this transient interaction and initiate signalling. Such a competition-based view of G protein versus DVL coupling at FZDs may explain why robust GTPγS binding was observed in response to Wnt5a in microglia membrane preparations, in which DVL is lost but G protein is retained^14^. In addition, as discussed below, the presence of lipids or a specific lipid composition may further facilitate G protein coupling and signalling.

The geometry of the intracellular cavity — hydrophobic grooves accommodating two acyl tails and flanked by positively charged residues that coordinate polar headgroups — is characteristic of a lipid-binding site. It is possible that in the cell, this site is occupied by an endogenous lipid. Lipids, particularly phosphoinositides, are established modulators of GPCR signalling^42^, and PIP2 has numerous roles in facilitating Wnt signalling. Wnt stimulation leads to PIP2 production to cause LRP5/6 phosphorylation^43–45^, aid in the recruitment of signalosome forming proteins^46,47^, enhance DEP domain affinity for FZD4^48^, and regulate complex formation between DEP domain and FZD6^49^. Moreover, within class F GPCRs, SMO is a lipid-binding GPCR that recruits Gi upon cholesterol binding. Whilst our lipidomic analyses identified a number of PIP lipid species enriched in GDN-purified samples relative to LMNG- or Quad-purified complexes, increasing cellular PIP levels did not detectably alter FZD-mediated G protein activation. However, these data do not conclusively exclude an endogenous lipid. The pocket might bind a diverse or substoichiometric mixture of lipids, where individual species would be averaged during cryo-EM processing, obscured by the high abundance of other lipid species in mass spectrometry or have high specificity for the identity of the lipid tails. It remains plausible that a specific endogenous lipid or lipid-like molecule binds to this site and exerts an effect in a cellular context.

Taken together, our findings reframe FZDs as inherently poor G protein activators whose weak coupling is a consequence of conserved structural features of the intracellular cavity, and instead predispose FZDs to DVL coupling and signalling. LMNG has served as an inadvertent tool that has made these FZD-G protein complexes tractable for structural study, but in doing so has obscured the dynamic nature of FZD–G protein engagement. Future studies aimed at capturing physiologically relevant FZD activation states will need to carefully consider both the choice of detergent and the potential influence of competing intracellular effectors.

## Methods

### Expression and purification of Nb35

Nb35^50^ was expressed in the periplasm of *E. coli* BL21(DE3) Rosetta cells cultured at 37 °C in LB broth containing 100 µg/mL carbenicillin and 35 µg/mL chloramphenicol. At OD₆₀₀ ∼0.7, expression was induced with 1 mM IPTG and temperature reduced to 18 °C for ∼16 hours. Cells were pelleted, resuspended in ice-cold lysis buffer (0.2 M Tris pH 8.0, 0.5 M sucrose, 0.5 mM EDTA; 5 mL/g pellet), and incubated for 1 hour at 4 °C. Two volumes of ice-cold MQ water were added and incubation continued for 45 minutes. Following centrifugation, the periplasmic supernatant was adjusted to 20 mM HEPES pH 7.5, 150 mM NaCl, 5 mM MgCl₂, and 5 mM imidazole, then batch-bound to Ni-NTA resin for 90 minutes at 4 °C. Resin was washed with 40 CV of wash buffer (20 mM HEPES pH 7.5, 500 mM NaCl, 5 mM imidazole) and eluted with 20 mM HEPES pH 7.5, 100 mM NaCl, 250 mM imidazole. Eluate was concentrated using a 10 kDa MWCO centrifugal filter (Millipore), flash-frozen in liquid nitrogen, and stored at −80 °C.

### FZD5/7 – G protein complex expression and purification

FZD7 receptor construct included an N-terminal HA signal sequence, anti-FLAG epitope tag followed by a 3C protease cleavage site, full length human FZD7 sequence followed by a C-terminal 3C cleavage site, GGGS linker, 8×His tag and a miniGα_s_ fusion^51,52^. For the FZD7-Quad variant, four mutations were introduced: E286^2.41^A, A365^3.54^F, K466^6.28^E, and L467^6.29^T. The FZD5 construct consisted of full length human FZD5 sequence with the same N-terminal modifications as used at FZD7, but with the C-terminal tags arranged as 8×His tag, GGGS linker, 3C cleavage site, and fused to a modified mGα_q_ chimeric construct that is a mini-Gα_s_ substituted with Gα_q_ residues at the receptor interface and the αN of Gα_i_^53,54^. The chimeric receptors were cloned into pACEBac1 (Geneva Biotech); Gβ_1_ (C-terminally 8×His-tagged) and Gγ_2_ were cloned into pVL1392 (Expression Systems). FZD7-mGα_s_/FZD5-mGα_sq_ and Gβ_1_γ_2_ were co-expressed in *Trichoplusia ni* (Hi5) insect cells using the Bac-to-Bac Baculovirus Expression System (Invitrogen). Cells were cultured in ESF 921 serum-free medium (Expression Systems), infected at 3–4 × 10⁶ cells/mL, and shaken at 27 °C for ∼62 hours. Harvested cell pellets were flash-frozen in liquid nitrogen and stored at −80 °C.

For purification, pellets were resuspended and lysed in 20 mM HEPES pH 8.0, 50 mM NaCl, 2 mM MgCl₂, benzonase, and protease inhibitors (500 µM PMSF, 1 mM leupeptin, 1 mM benzamidine). For FZD7 only, 200 uM SAG1.3 was included. Nb35 and apyrase were added to promote complex formation, and the lysate was stirred at room temperature for 1 hour. Following centrifugation, the membrane pellet was solubilised for 90 minutes at 4 °C in 20 mM HEPES pH 8.0, 100 mM NaCl, 0.5% LMNG, 0.05% cholesterol hemisuccinate (CHS), 10% glycerol, 2 mM MgCl₂, 2 mM CaCl₂, benzonase, and protease inhibitors. The clarified supernatant was incubated with M1 anti-FLAG affinity resin for 1 hour at 4 °C, loaded onto a gravity column, and washed with 20 mM HEPES pH 8.0, 100 mM NaCl, 0.01% LMNG, 0.001% CHS, 2 mM MgCl₂, 2 mM CaCl₂. Protein was eluted in wash buffer lacking CaCl₂ and supplemented with 10 mM EGTA and 0.1 mg/mL FLAG peptide. The eluate was concentrated and subjected to size-exclusion chromatography (SEC) on a Superdex 200 Increase 10/300 column (GE Healthcare) in 20 mM HEPES pH 8.0, 100 mM NaCl, 0.01% LMNG, 0.001% CHS, 2 mM MgCl₂. Peak fractions were concentrated, flash-frozen in liquid nitrogen, and stored at −80 °C.

For FZD7-mGs complex purification in n-dodecyl-β-D-maltoside (DDM), the same protocol was used with the following modifications. Solubilisation buffer: 20 mM HEPES pH 8.0, 100 mM NaCl, 1% DDM, 0.05% CHS, 10% glycerol, 2 mM MgCl_2_, 2 mM CaCl_2_, protease inhibitors (500 μM PMSF, 1 mM leupeptin, 1 mM benzamidine) and benzonase. Wash buffer: 20 mM HEPES pH 8.0, 100 mM NaCl, 0.1% DDM, 0.005% CHS, 2 mM MgCl_2_, 2 mM CaCl_2_. FLAG elution: 20 mM HEPES pH 8.0, 100 mM NaCl, 0.1% DDM, 0.005% CHS, 2 mM MgCl_2_, 10 mM EGTA, 0.1 mg/mL FLAG peptide. Size exclusion buffer: 20 mM HEPES pH 8.0, 100 mM NaCl, 0.01% LMNG, 0.001% CHS, 2 mM MgCl_2._

For FZD7-mGs complex purification in glyco-diosgenin (GDN), the same protocol was used with the following modifications. Solubilisation buffer: 20 mM HEPES pH 8.0, 100 mM NaCl, 1% DDM, 0.05% CHS, 10% glycerol, 2 mM MgCl_2_, 2 mM CaCl_2_, protease inhibitors (500 μM PMSF, 1 mM leupeptin, 1 mM benzamidine) and benzonase. Protein bound FLAG column was washed with 20 column volumes (CV) of DDM wash buffer (20 mM HEPES pH 8.0, 100 mM NaCl, 0.1% DDM, 0.005% CHS, 2 mM MgCl_2_, 2 mM CaCl_2_). Detergent was gradually exchanged to GDN with 6 different detergent mixes (50:50, 75:25, 90:10, 95:5, 99:1, 100:0) of DDM wash buffer to GDN wash buffer (20 mM HEPES pH 8.0, 100 mM NaCl, 0.1% GDN, 0.01% CHS, 2 mM MgCl_2_, 2 mM CaCl_2_). Protein was subsequently washed with 20 mM HEPES pH 8.0, 100 mM NaCl, 0.0525% GDN, 0.00525% CHS, 2 mM MgCl_2_, 2 mM CaCl_2_ followed by 20 mM HEPES pH 8.0, 100 mM NaCl, 0.021% GDN, 0.0021% CHS, 2 mM MgCl_2_, 2 mM CaCl_2._ Protein was eluted using 20 mM HEPES pH 8.0, 100 mM NaCl, 0.01% GDN, 0.001% CHS, 2 mM MgCl_2_, 10 mM EGTA, 0.1 mg/mL FLAG peptide. Size exclusion buffer: 20 mM HEPES pH 8.0, 100 mM NaCl, 0.0021% GDN, 0.00021% CHS, 2 mM MgCl_2_.

### Membrane scaffold protein expression and purification

MSP1D1 (Addgene #20061)^55^ was expressed in *E. coli* BL21(DE3) cells grown in S broth with 50 µg/mL kanamycin. Expression was induced at OD₆₀₀ ∼0.7 with 1 mM IPTG for 3 hours, after which cells were pelleted and stored at −80 °C. Pellets were resuspended in lysis buffer (50 mM Tris-HCl pH 8.0, 250 mM NaCl, 0.5% Triton X-100, 0.5 mM EDTA, 5 mM MgCl₂, protease inhibitor cocktail (Roche), benzonase (Sigma)) with lysozyme added to 1 mg/mL. After 30 minutes at room temperature, lysate was sonicated and clarified by centrifugation. The supernatant was supplemented with 5 mM imidazole and batch-bound to Ni-NTA resin for 1 hour at 4 °C. Resin was loaded onto a gravity column and washed sequentially with 10 CV each of: (a) 40 mM Tris-HCl pH 8.0, 300 mM NaCl, 1% Triton X-100; (b) 40 mM Tris-HCl pH 8.0, 300 mM NaCl, 50 mM sodium cholate, 20 mM imidazole; and (c) 40 mM Tris-HCl pH 8.0, 300 mM NaCl, 20 mM sodium cholate, 40 mM imidazole. Protein was eluted in 40 mM Tris-HCl pH 8.0, 300 mM NaCl, 5 mM sodium cholate, 300 mM imidazole, then dialysed overnight at 4 °C into 10 mM Tris-HCl pH 7.4, 25 mM NaCl, 1 mM EDTA. MSP1D1 was further purified by anion exchange chromatography on a HiTrap Q FF column (Cytiva) equilibrated in buffer A (10 mM Tris-HCl pH 7.4, 25 mM NaCl, 1 mM EDTA). After washing with 3 CV of buffer A, protein was eluted over a 30 CV linear gradient to buffer B (10 mM Tris-HCl pH 7.4, 300 mM NaCl, 1 mM EDTA). The eluate was dialysed into 10 mM Tris-HCl pH 7.4, 100 mM NaCl, 1 mM EDTA, concentrated using a 10 kDa MWCO centrifugal filter, flash-frozen in liquid nitrogen, and stored at −80 °C.

### FZD7-mGs nanodisc reconstitution

POPC and POPG (Avanti Polar Lipids) were dissolved in chloroform at 10 mM and 6.67 mM respectively (3:2 molar ratio), dried under a nitrogen stream to form a thin film, and resuspended in HNE buffer (20 mM HEPES pH 8.0, 100 mM NaCl, 1 mM EDTA) supplemented with 50 mM sodium cholate. For nanodisc reconstitution, 5 µM DDM purified FZD7-mGs complex, 100 µM MSP1D1, and 5 mM POPC:POPG were mixed in 730 µL HNE buffer and incubated on ice for 1 hour. Detergent was removed by sequential incubation with Bio-Beads SM-2 (Bio-Rad): twice with 50 mg for 1 hour each, then overnight with 200 mg, all at 4 °C with rotation. Bio-Beads were removed by filtration and rinsed with 400 µL HNE buffer, which was pooled with the main sample. The nanodisc-incorporated complex was purified by supplementing the sample with 2 mM MgCl₂ and incubating with M1 anti-FLAG affinity resin for 1 hour at 4 °C. Resin was loaded onto a gravity column, washed with 25 CV of HNE buffer containing 2 mM MgCl₂, and eluted in HNE buffer + 10 mM EGTA and 0.1 mg/mL FLAG peptide. The eluate was concentrated and subjected to SEC on a Superdex 200 Increase 10/300 GL column (GE Healthcare) equilibrated in HNE buffer. Peak fractions were pooled, concentrated using a 100 kDa MWCO centrifugal filter (Millipore), flash-frozen in liquid nitrogen, and stored at −80 °C.

### Vitrified sample preparation and data collection

For all FZD-G protein complexes, 3 µL of sample was applied to glow-discharged UltrAuFoil R1.2/1.3 Au 300 mesh grids (Quantifoil). Protein concentrations of samples were as follows. WT FZD5-mGq LMNG: 15 AU, WT FZD7-mGs LMNG: 15 AU, WT FZD7-mGs Nanodisc: 5 AU, WT FZD7-mGs GDN: 22 AU and Quad Mutant FZD7-mGs LMNG: 5 AU. Grids were vitrified using a Vitrobot Mark IV (Thermo Fisher Scientific) operated at 4 °C and 100% humidity, with a blot time of 5 s and a blot force of 15. Imaging was carried out on a Titan Krios microscope operating at an accelerating voltage of 300 kV in nanoprobe EFTEM mode, using a 50 µm C2 aperture and a 100 µm objective aperture, with zero-loss filtering at a 10 eV slit width. Data were collected with aberration-free image shift (AFIS) in EPU (Thermo Fisher Scientific). Additional acquisition parameters for each dataset are given in Supplementary Table 1.

### Image Processing

Specific details for the processing of all FZD-G protein complexes are found in Supplementary Fig. 3-6,9. All movies were processed by correcting for beam-induced motion using Patch Motion correction, followed by Patch CTF Estimation in cryoSPARC v4.7.0^56^. Micrographs with an estimated CTF fit resolution worse than 4 Å were discarded. All resolutions are reported according to the gold-standard Fourier shell correlation (FSC) criterion at 0.143, and directional resolution anisotropy was assessed from directional FSC (3DFSC) plots and the corresponding viewing-direction distributions.

### WT FZD5-mGq complex in LMNG

We collected 2 datasets comprising 4,623 and 6,750 movies, respectively, at 0.82 Å/pix. Each was processed independently before combining the best particles for a final structure. All movies were subjected to patch motion correction and patch CTF refinement. A small subset of micrographs was used to generate an initial model, which was subsequently used for template picking. Template picking of each dataset yielded 3.9 and 5.8 million particles, respectively. These were extracted (4x binned, 64 pix, 3.28 Å/pix), and junk particles were removed through three rounds of heterogeneous refinement in cryoSPARC (with a junk model included as a decoy class). The datasets were further cleaned using 2D classification. Particles were then re-extracted at 0.82 Å/pixel in a 256-pixel box. At this point, the maps appeared extremely anisotropic, with a strong preferred orientation, and were refractory to any particle-rebalancing attempts. Instead, the dataset was imported into RELION 4.0^57^ and subjected to 3D classification. Particles belonging to the best class were re-imported into cryoSPARC and subjected to non-uniform refinement and local motion correction. At this point, the datasets — containing 203,677 and 273,343 particles, respectively — were combined for a final round of non-uniform refinement. The final global resolution was 2.72 Å (0.143 FSC criterion).

### WT FZD7-mGs complex in LMNG

6,444 movies at 0.82 Å/pixel were collected with 5,972 micrographs after micrograph processing. Particles were selected by template picking, using templates generated from a subset of the data, yielding 8,630,449 picks. 8,214,533 particles were extracted in an 80 pixel box with 4x binning (3.28 Å/pixel). Three rounds of heterogeneous refinement, performed against a map generated from a subset of the data together with a junk class as a decoy generated 2,747,017 particles. 2D classification into 100 classes gave 2,029,885 particles, which were re-extracted unbinned in a 320-pixel box (0.82 Å/pixel). 2D classification into 100 classes retained 1,684,381 particles. These were subjected to two rounds of heterogeneous refinement against ab initio references (948,206 particles), followed by a final round of 2D classification into 100 classes to give 832,537 particles. This final particle set was refined by non-uniform refinement, subjected to local (reference-based) motion correction, and refined again by non-uniform refinement, producing a final consensus map (2.61 Å). A further local refinement was performed to generate a receptor-focused map (2.53 Å).

### WT FZD7-mGs complex in lipid nanodisc

11,217 movies at 0.82 Å/pixel were collected with 9,393 micrographs after micrograph processing. Particles were selected by template picking, yielding 10,695,632 picks, of which 9,316,487 particles were extracted in an 80-pixel box with 4x binning (3.28 Å/pixel). To generate suitable 3D references, a subset of 3,197,898 particles obtained from two rounds of 2D classification into 200 classes was put through an iterative procedure of heterogeneous refinement, 2D classification and ab-initio reconstruction, producing three volumes representing the complex together with a junk class. The full extracted particle stack was then subjected to three rounds of heterogeneous refinement against these references, reducing the stack to 1,539,336 particles. Two rounds of 2D classification into 200 classes gave 697,943 particles that were re-extracted unbinned in a 320-pixel box (0.82 Å/pixel). This particle set was refined by non-uniform refinement, subjected to reference-based motion correction, and refined again by non-uniform refinement. Subsequent 3D classification separated the particles into three classes of 360,099 particles, 310,380 particles and 24,292 particles. The two classes showing well-resolved density for the complex (670,479 particles in total) were combined and refined by non-uniform refinement to give a final consensus map (2.37 Å). A further local refinement was performed to generate a receptor-focused map (2.78 Å).

### WT FZD7-mGs complex in GDN micelle

21,181 movies at 0.82 Å/pixel were collected with 20,585 micrographs after micrograph processing. Particles were selected by template picking, using templates generated from a subset of the data, yielding 22,983,065 picks. These were sorted by four rounds of heterogeneous refinement against a set of references comprising maps generated from a subset of the data together with a junk class, retaining 1,015,012 particles in the classes showing protein features. These particles were re-extracted with 2x binning in a 180-pixel box (1.64 Å/pixel) and subjected to a further round of heterogeneous refinement against maps generated from a subset of the data and a junk class, giving 611,332 particles. Ab initio reconstruction of this particle set produced one volume corresponding to the complex and three junk classes, and a further round of heterogeneous refinement of the same particles against maps generated from a subset of the data together with these ab initio junk classes gave 387,672 particles. Particles were then re-extracted unbinned in a 360-pixel box (0.82 Å/pixel) and refined by non-uniform refinement, reference-based motion correction, and further non-uniform refinement leaving 386,319 particles. To further improve homogeneity, two rounds of 2D classification into 100 classes were used to select 131,698 particles, from which ab initio reconstruction generated one volume of the complex and two junk classes. The 386,319-particle set was then classified by three rounds of heterogeneous refinement against these references, and the resulting 78,395 particles were refined by non-uniform refinement to give the final consensus map (3.02 Å).

### Quad Mutant FZD7-mGs complex in LMNG micelle

9,890 movies at 0.82 Å/pixel were collected with 7,439 micrographs after micrograph processing. Template picking, using templates generated from a subset of the data, yielded 9,464,616 picks that were extracted in an 80-pixel box with 4x binning (3.28 Å/pixel). Three rounds of heterogeneous refinement against an ab initio map together with a junk class retained 2,244,028 particles. A subset of 200,000 particles was used for ab initio reconstruction to generate five references. All 2,244,028 particles were then classified against these references by three rounds of heterogeneous refinement, retaining 289,059 particles. A round of 2D classification into 200 classes (256,272 particles) was followed by a further 3D heterogeneous refinement (194,652 particles). These particles were re-extracted unbinned in a 320-pixel box (0.82 Å/pixel) giving 188,715 particles, which were refined by non-uniform refinement. A further round of 3D heterogeneous refinement gave 107,883 particles. Following reference-based motion correction, 2D classification into 100 classes retained 102,360 particles, and a final 3D heterogeneous refinement gave 78,093 particles. Non-uniform refinement of this set produced the final consensus map (2.96 Å). A further local refinement was performed to generate a receptor-focused map (3.30 Å).

### Model building and refinement

The starting model for FZD7-mGs was derived from PDB entry 7EVW whilst the starting model for FZD5-mGq was generated using AlphaFold2^58^. Models were docked into the EM maps in UCSF ChimeraX^59^ and subjected to rigid-body fitting in PHENIX^60^. Refinement proceeded through iterative cycles of manual building in Coot^61^ and ISOLDE^62^, with real-space refinement in PHENIX. Restraints for LMNG were obtained from the GRADE web server (https://grade.globalphasing.org). Models were validated using MolProbity^63^ and the wwPDB validation server^64^. Figures were prepared in UCSF ChimeraX and PyMOL (Schrödinger). Surface area calculations were performed through use of QtPISA v2.1.0^65,66^.

### Molecular Dynamics simulations

All molecular dynamics (MD) simulations of the membrane-embedded FZD7 complex were performed using GROMACS 2023^67–69^ in combination with the GROMOS 54a7 force field^70^. Several regions of the cryo-EM structure lacked sufficient electron density for reliable atomic modelling and were therefore excluded. Within the Gα subunit, this applied to N-terminal residues 1-8 and residues 254-264. Additionally, because the structure was resolved using an mGα_s_ construct, the α-helical domain (residues 60-205) was absent from the model. For the Gβ subunit, residues 1-3 at the N-terminus and residues 129-131 could not be modelled. For the Gγ subunit, N-terminal residues 1-7 and C-terminal residues 64-71 were unresolved and unmodelled. No density was observed for the N-terminal domain of FZD7 and as such residues 1-205 were not modelled. Similarly for the C-terminal domain residues 562-574. The same residues were missing and unmodelled for the FZD7-Quad-mGs complex where Nb35 was also removed from the model. For all these residues, acetyl and NH_2_ caps were applied to avoid the introduction of inappropriate charges. Parameters for LMNG were generated using the Automated Topology Builder 4.0 (molid 1834487)^71^.

Four simulation systems were prepared: (1) the FZD7-mGs complex with LMNG bound, (2) the FZD7-mGs complex with LMNG removed, (3) the FZD7-Quad-mGs complex, and (4) an FZD7 receptor-only system containing LMNG, generated by removal of the G protein heterotrimer from the FZD7-mGs complex. All systems were embedded in a lipid bilayer composed of 80% POPC and 20% cholesterol, constructed using the MemGen webserver^72^. Each system was subsequently solvated with simple point charge (SPC) water molecules, 150 mM NaCl was added to the solvent to approximate physiological ionic conditions^73^. Counterions were added to ensure overall charge neutrality of the system. All simulations were performed under periodic boundary conditions with hydrogen mass repartitioning applied^74^, with the Silico package used for LMNG^75^. Each system underwent energy minimisation via the steepest descent algorithm, followed by a 10 ns stepwise equilibration using a 2 fs timestep. During the equilibration, harmonic position restraints of 1000 kJ/mol/nm^2^ were initially applied to the protein and ligand atoms. These restraints were progressively reduced in 2 ns time intervals to 500 and 100 kJ/mol/nm^2^, followed by a final 4 ns of equilibration with position restraints of 10 kJ/mol/nm^2^.

The coordinates of the final frame of each equilibrated system were used as the starting configuration for production simulations. Production simulations were performed in the NPT ensemble. Temperature was maintained at 310 K using the Bussi-Donadio-Parrinello velocity-rescaling thermostat^76^ with a coupling time constant of 0.1 ps, applied separately to solvent and solute groups. Initial velocities were assigned from a Boltzmann distribution at 310 K. Semi-isotropic pressure was maintained at 1.0 bar using the Parrinello-Rahman barostat^77^ with a coupling time constant of 40 ps and a compressibility of 4.5 x 10⁻⁵ bar⁻¹. The geometry of water molecules was constrained using the SETTLE algorithm^78^, while covalent bond lengths within the solute were constrained using LINCS^79^. Long-range electrostatic interactions were calculated using Particle Mesh Ewald (PME) summation. Non-bonded interactions were evaluated using the Verlet cutoff scheme with a cutoff distance of 1.4 nm. Three independent replicate simulations of 500 ns were performed for each system, with randomised initial velocities. Simulations were visualised using VMD v1.9.3^80^ and analysed using GROMACS tools.

### Expression and purification of Wnt3a

Expi293 (ThermoFisher Scientific) cells grown in adherent cultures in Dulbecco’s Modified Eagle Medium (DMEM) High glucose, GlutaMAX (ThermoFisher), 10% Fetal Bovine Serum (FBS) in 5% CO_2_, 37 °C were stably transfected with His-Wnt3a (human Wnt3a, carrying an N-terminal 6xHis between its native signal sequence and a mature protein sequence). Conditioned media (C.M.) was collected and incubated with TALON resin overnight at 4 °C with rotation. Resin was washed with 20 mM HEPES, 150 mM NaCl, pH 8.0 and eluted using the same buffer supplemented with 250 mM imidazole. Protein was dialysed in 20 mM HEPES, 150 mM NaCl, pH 8.0 and concentrated. To determine the concentration of active Wnt3a within the protein sample, titrating volumes of the protein elution were tested against known amounts of commercial Wnt3a (R&D Systems, 5036-WN) using the TOPFlash assay (described below).

### Mammalian cell culture

Human Embryonic Kidney (HEK) 293T cells were grown in DMEM supplemented with 2 mM GlutaMAX (ThermoFisher) and 10% FBS in a humidified incubator at 37°C and 5% CO_2_. Once cells reached 60-80% confluency, the medium was aspirated, and the cells were washed with phosphate-buffered saline (PBS). Cells were detached using Versene (0.48 mM EDTA in PBS), collected, and centrifuged at 350 × g for 3 minutes. The cell pellet was resuspended, and cells were either plated for subsequent assays or passaged into new tissue culture flasks for continued growth.

### BRET GDPβS Constitutive Activity Assay

ΔFZD1-10 HEK293T^81^ cells were seeded in 6-well plates at 2 × 10⁵ cells/mL (2 mL per well) 24 hours prior to transfection. Adherent cells were transfected using polyethylenimine (PEI; 1:4 DNA:PEI ratio) with a plasmid mixture comprising 250 ng receptor–nLuc, 750 ng Gα_s_, 250 ng Gβ_1_, and 250 ng Gγ_2_–Venus. To suppress endogenous Wnt ligand activity, cells were treated with 100 nM C59 concurrent with transfection. 24 hours post-transfection, cells were detached using Versene, and transferred to 96-well white-walled CELLSTAR plates (Greiner Bio-One) at 30,000 cells per well for a further 24 hours. On the day of the assay, growth medium was aspirated and wells washed with 200 µL phosphate-buffered saline (PBS). Cells were then resuspended in 80 µL assay buffer (140 mM KCl, 10 mM NaCl, 1 mM MgCl₂, 0.1 mM EGTA, 20 mM HEPES, 10 µg/mL digitonin, 2 U/mL apyrase). Cells were incubated at 37 °C for 30 minutes. Bioluminescence resonance energy transfer (BRET) was initiated by addition of 10 µL NanoGlo luciferase substrate (1:1000 final dilution). Following a 10-minute equilibration period, four baseline BRET readings were acquired using a Clariostar plate reader (BMG Labtech), measuring nLuc donor emission at 465 ± 20 nm and Venus acceptor emission at 540 ± 20 nm. Cells were then stimulated by addition of 10 µL of either assay buffer (vehicle) or GDPβS (Sigma-Aldrich, G7637; 10 µM final concentration). BRET was recorded continuously for 30 minutes following stimulation. BRET signal was calculated as the ratio of acceptor emission over donor emission. The ratio was well-corrected using the first four baseline measurements to give ΔBRET ratio and then corrected again using the vehicle treated wells to give ΔΔBRET ratio. All data were analysed using GraphPad Prism 10.

### TRUPATH

ΔFZD1-10 HEK293T cells were transfected transiently in suspension using a 1:4 ratio DNA to PEI. Each transfection contained 10 ng per well of the G protein TRUPATH biosensors^33^ pcDNA5/FRT/TO-Gα_sS/q/i1_-rLuc8/pcDNA3.1-Gβ_3_/pcDNA3.1-Gγ_9_-GFP2 and 0–50 ng per well of either WT FZD5/7, or mutant FZD7 constructs (E286^2.41^A, A365^3.54^F, K466^6.28^E, L467^6.29^T and E286^2.41^A/A365^3.54^F/K466^6.28^E/L467^6.29^T). The total DNA per well was adjusted to 80 ng using empty pcDNA3.1 vector. Cells were seeded at a density of 30,000 cells per well into white-walled 96-well CELLSTAR plates (Greiner Bio-One). After 48 hours, cells were washed with 200 μL PBS and replaced with 1x Hanks’ Balanced Salt Solution (HBSS) supplemented with 10 mM HEPES pH 7.4. Plates were incubated at 37°C for 30 minutes and 10 μL of 1.3 μM Prolume Purple coelenterazine (NanoLight Technology, Pinetop, AZ) was added to each well. Cells were incubated for an additional 10 minutes at 37°C prior to BRET detection. BRET emission was recorded over fourteen consecutive 100-second intervals using a PHERAstar FSX plate reader (BMG Labtech) with 410 ± 80 nm (donor) and 515 ± 30 nm (acceptor) emission filters. BRET signal was calculated as the ratio of 515 ± 30-nm emission over 410 ± 80-nm emission. The ratio was baseline corrected using the 0 ng receptor condition to give ΔBRET ratio. All data were analysed using GraphPad Prism 10.

To determine the influence of lipids on constitutive activity, ΔFZD1-10 HEK293T cells were transfected and seeded as described above. 18:1 PI(3)P1, 18:1 PI(4)P1,18:1 PI(5)P1, C18:1 LPA, 18:1 PI(3,4)P2, 18:1 PI(3,5)P2, 18:1 PI(4,5)P2 (Avanti Lipids) were dissolved in water:methanol:chloroform (1:9:20), dried under a stream of nitrogen gas and resuspended to 500 µM in 20 mM HEPES pH 7.4, 100 mM NaCl. Unlabelled Shuttle PIP Carrier 2 (PIP2) and Unlabelled Shuttle PIP Carrier 3 (LPA and PIP1, Echelon Biosciences) were resuspended to 500 µM in the same buffer. Lipid and carrier were mixed 1:1 (final concentration of 250 µM each) and incubated at RT for 10 minutes; for the carrier-only control, carrier was mixed 1:1 with buffer alone. Complexed lipids were diluted 10-fold in 1x HBSS supplemented with 10 mM HEPES pH 7.4 to a final concentration of 25 µM. At 48 hours post-transfection, cells were washed with 200 µL PBS and 100 µL of lipid solution was added per well. Plates were incubated at 37°C for 30 minutes. BRET detection and analysis were carried out as described above.

### CAMYEL

ΔFZD1-10 HEK293T or ΔDVL1-3 HEK293T cells were transfected transiently in suspension using a 1:4 ratio DNA to PEI. Each transfection contained 20 ng per well of the cAMP CAMYEL biosensor^82^ (ATCC, Manassas, VA, USA) alongside titrated amounts of WT FZD7 or FZD7-Quad plasmid (0, 2.5, 5, 10, or 50 ng/well). The total DNA per well was adjusted to 70 ng using empty pcDNA3.1 vector. To suppress endogenous Wnt secretion, cells were treated with 100 nM C59 (a Porcupine inhibitor) concurrent with transfection. Forty-eight hours post-transfection, cells were washed once with 200 µL PBS and equilibrated in HBSS supplemented with 10 mM HEPES (pH 7.4) for 30 minutes at 37 °C. Coelenterazine-H (Nanolight Technology; dissolved to a 3.3 mM stock in manufacturer-supplied solvent) was added to a final concentration of 5 µM and incubated for 10 minutes at 37 °C prior to BRET measurement. For the 0 ng/well condition, 100 µM forskolin was included as a positive control for cAMP elevation, added concurrently with coelenterazine-H. BRET was recorded for 30 minutes on a CLARIOstar plate reader (BMG Labtech), with Renilla luciferase (rLuc) donor emission detected at 475 ± 15 nm and YFP acceptor emission detected at 535 ± 15 nm. BRET signal was calculated as the ratio of 535 ± 15 nm emission over 475 ± 15 nm emission. The ratio was baseline corrected using the 0 ng receptor condition to give ΔBRET ratio which was subsequently normalise to zero and the response observed for 100 µM forskolin. All data were analysed using GraphPad Prism 10.

### Flow cytometry

To determine the expression of WT FZD7 and mutants in the TRUPATH and CAMYEL assays in parallel, transfected cells were harvested by incubating with 100 μL of Versene for 5–10 minutes at 37 °C to facilitate detachment, followed by transfer to an ice-cold 96-well U-bottom plate. Cells were pelleted by centrifugation at 350 × g for 5 minutes at 4 °C, then resuspended in blocking buffer (1× PBS, 1 mM EDTA, 2 mM CaCl₂, 5% BSA) and incubated for 45 minutes on ice. After blocking, cells were centrifuged again at 350 × g for 5 minutes, resuspended in 100 μL FACS buffer (1× PBS, 1 mM EDTA, 2 mM CaCl₂, 0.1% BSA) containing 333 ng/mL of AF700 conjugated M1 anti-FLAG antibody and stained for 1 hour on ice. Following extracellular staining, cells were fixed and permeabilised through use of BD Cytofix/Cytoperm Fixation Permeabilisation Kit (BD Biosciences). Cells were washed twice with FACS buffer then permeabilised by resuspension in 100 μL of fixation/permeabilization solution for 20 minutes on ice. Cells were washed twice with diluted Perm/Wash buffer (1:9 dilution in MQ), with a 15-minute incubation during the first wash step to aid blocking, and centrifuged at 350 × g for 5 minutes. Intracellular staining was performed by resuspending cells in 100 μL of Perm/Wash buffer containing AF700-conjugated anti-FLAG antibody (at 333 ng/mL) and 2 mM CaCl₂, followed by a 30-minute incubation at room temperature in the dark. After staining, cells were washed twice and resuspended in Perm/Wash buffer for acquisition on a NovoCyte Quanteon Flow Cytometer. All data was analysed using FlowJo v10 software.

### Saturation net BRET

ΔFZD1-10 HEK293T or ΔDVL1-3 HEK293T cells were seeded at a density of 30,000 cells per well in white-walled 96-well CELLSTAR plates (Greiner Bio-One) and transiently transfected in suspension with 20 ng per well of receptor–NanoLuc (nLuc) construct and increasing amounts of Venus–DVL2^83^, mVenus-mGα_s_^84^ or Venus-WT Gα_s_ plasmid to achieve nLuc:Venus DNA ratios ranging from 1:0 to 1:10. All wells were adjusted to a total of 220 ng DNA using empty pcDNA3.1 vector. Cells were treated with 100 nM C59 (a Porcupine inhibitor) to suppress endogenous Wnt ligand activity. After 48 hours, media were aspirated, and cells were washed with 200 μL of pre-warmed HBSS supplemented with 10 mM HEPES. Cells were then incubated in 90 μL of the same buffer at 37 °C for 20–30 minutes prior to detection. First, Venus fluorescence intensity was measured using a CLARIOstar plate reader (BMG Labtech) with excitation at 485 ± 10 nm and emission at 535 ± 10 nm. Subsequently, 10 μL of furimazine substrate (Promega Nano-Glo Luciferase Assay System, N1110) prepared in HBSS + 10 mM HEPES was added to each well (final dilution 1:1000), followed by a 10-minute incubation at 37 °C. Four baseline BRET measurements were acquired with nLuc emission detected at 465 ± 20 nm and Venus emission at 540 ± 20 nm. Ten additional BRET readings were then recorded. The initial Venus fluorescence intensity (pre-furimazine) values were divided by the first four nLuc luminescence readings (post-furimazine) to give a Venus fluorescence/nLuc luminescence ratio for each receptor:mVenus–DVL2/Venus-Gs ratio. All Venus fluorescence/nLuc luminescence ratios were background-corrected with the 1:0 control ratio. Net BRET values were calculated by subtracting the background BRET observed at the 1:0 from the average BRET signal obtained from the final ten BRET measurements. Saturation binding curves were generated by plotting the corrected fluorescence to luminescence ratios against the corresponding net BRET values using GraphPad Prism 10.

### TOPFlash

ΔFZD1-10 HEK293T cells were transfected transiently in suspension using a 1:4 ratio of DNA to PEI with 10 ng per well of WT FZD7 or mutants, 20 ng per well of Super8xTopFlash^85^ and 4 ng per well of pRL-TK Renilla (34 ng of DNA in total). Cells were plated at 40,000 cells per well into clear F-bottom 96-well microplates precoated with 50 μL of 1 mg/mL poly-D-lysine. After 24 hours, media was replaced with fresh media containing Wnt3a. Twenty-four hours post-stimulation, cells were washed with phosphate-buffered saline (PBS), and luciferase activity was measured using the Dual-Luciferase® Reporter Assay System (Promega). Cells were lysed in 15 μL of Passive Lysis Buffer with shaking at 4 °C for 2 hours. Lysates were transferred to a white half-area 96-well microplate (Greiner Bio-One) for luminescence measurements. Firefly and Renilla luciferase signals were sequentially measured using a CLARIOstar plate reader (BMG Labtech). For each well, 25 μL of Luciferase Assay Reagent II was injected to initiate the firefly signal, which was recorded at 580 ± 40 nm over ten intervals of 1.3 seconds each. Subsequently, 25 μL of Stop & Glo reagent was injected to quench firefly activity and initiate Renilla luminescence, which was detected at 480 ± 40 nm under the same acquisition settings. The TOPFlash signal was calculated by dividing the AUC for firefly (580 ± 40 nm) emission by the AUC for Renilla (480 ± 40 nm) emission. Data were normalised to values from buffer only and 3000 ng/mL Wnt3a conditions at WT FZD7. Concentration response curves were fit using a non-linear four parameter logistic equation in GraphPad Prism 10 to derive E_max_ and Log EC_50_ values.

### LC–QTOF-MS screening of detergents in nanodiscs and isolated membrane-protein complexes

#### Initial screen

Liquid chromatography was performed using an Agilent LC system equipped with a binary pump, multisampler and thermostatted column compartment. Samples (1 µL) were injected, and the autosampler needle was washed for 12 s at the flush port following each injection using 50:30:20 isopropanol/acetonitrile/water (v/v/v).

Chromatographic separation was performed on an Agilent ZORBAX RRHD Eclipse Plus C18 column (2.1 x 100 mm, 1.8 µm) maintained at 45 °C, at a flow rate of 0.45 mL/min. Mobile phase A consisted of 50:30:20 water/acetonitrile/isopropanol (v/v/v) containing 10 mM ammonium formate and 5 µM medronic acid, while mobile phase B consisted of 1:9:90 water/acetonitrile/isopropanol (v/v/v) containing 10 mM ammonium formate. The initial composition of 90% A/10% B was maintained for 0.5 min, after which mobile phase B was increased linearly to 100% at 7.0 min and maintained until 9.0 min. The gradient was returned to 90% A/10% B by 9.5 min and maintained at the initial composition until the end of the 11-min run.

Mass spectrometric analysis was performed using an Agilent Revident Q-TOF mass spectrometer equipped with a dual Agilent Jet Stream electrospray ionisation source. The instrument was operated in positive-ion mode using the 1700 fragile/high-resolution acquisition configuration. Source conditions were: drying-gas temperature, 250 °C; drying-gas flow, 8 L/min; nebuliser pressure, 30 psi; sheath-gas temperature, 325 °C; sheath-gas flow, 11 L/min; capillary voltage, 4500 V; nozzle voltage, 500 V; fragmentor voltage, 175 V; skimmer voltage, 45 V; and octopole RF peak voltage, 750 V. Full-scan MS data were acquired over *m/z* 50–1700. The LC effluent was diverted to waste from 0–1 min, directed to the mass spectrometer from 1–10 min, and returned to waste after 10 min. Continuous online mass correction was performed using reference ions at *m/z* 121.050873 and 922.009798.

### Subsequent screens

An updated LC-QTOF-MS method was used for subsequent detergent screening. Chromatographic separation was performed using an Agilent LC system equipped with a binary pump, multisampler and thermostatted column compartment and an Agilent ZORBAX RRHD Eclipse Plus C18 column (2.1 x 100 mm, 1.8 µm) maintained at 55 °C. Samples (5 µL) were injected at a flow rate of 0.40 mL/min, with the autosampler needle and seat washed for 10 s following each injection.

Mobile phase A consisted of 50% methanol in water containing 5 mM ammonium formate, 2.5 µM medronic acid and 1 mM formic acid, while mobile phase B consisted of isopropanol. The gradient commenced at 100% A and was maintained until 0.5 min. Mobile phase B was then increased linearly to 95% at 8.0 min and maintained at 95% B until 10.0 min. The gradient was returned to 100% A by 10.5 min and maintained under the initial conditions until the end of the 12-min run.

Mass spectrometric analysis was performed using an Agilent Revident Q-TOF mass spectrometer equipped with a dual Agilent Jet Stream electrospray ionisation source and operated in positive-ion mode using the 1700 fragile/high-resolution acquisition configuration. Source conditions were: drying-gas temperature, 250 °C; drying-gas flow, 8 L/min; nebuliser pressure, 30 psi; sheath-gas temperature, 300 °C; sheath-gas flow, 11 L/min; capillary voltage, 4500 V; nozzle voltage, 1000 V; fragmentor voltage, 175 V; skimmer voltage, 45 V; and octopole RF peak voltage, 750 V.

### Lipidomic characterisation of purified membrane-protein complexes

#### Lipid extraction

Lipids were extracted using a single-phase extraction protocol modified from our previously described method^86^. Briefly, 100 µL of butanol/methanol/water (2:2:1, v/v/v) containing 1 mM ammonium formate and 50 µM medronic acid was added to each sample. A panel of stable isotope-labelled lipid internal standards was added, together with 40 pmol each of PI(3,5)P₂ 16:0/16:0-d5 and PI(5)P 16:0/16:0-d5 (Avanti Polar Lipids). Samples were vortex-mixed and sonicated in a water bath for 1 h, followed by centrifugation at 13,000 x g for 10 min. The resulting supernatant was transferred to glass autosampler vials for LC-MS analysis.

#### General lipidomics

Targeted lipidomic analysis was performed using an Agilent 1290 Infinity II LC system coupled to an Agilent 6495C triple-quadrupole mass spectrometer equipped with an Agilent Jet Stream electrospray ionisation source. Samples (5 µL) were injected, and the autosampler needle was washed at the flush port for 12 s following each injection. Chromatographic separation was performed on an Agilent Altura ZORBAX Eclipse Plus C18 column (2.1 x 100 mm, 1.8 µm) maintained at 45 °C at a flow rate of 0.40 mL/min.

The mobile-phase composition was based on our previously described C18 lipidomics methods^86,87^. Mobile phase A consisted of 50:30:20 water/acetonitrile/isopropanol (v/v/v) containing 10 mM ammonium formate and 5 µM medronic acid, while mobile phase B consisted of 1:9:90 water/acetonitrile/isopropanol (v/v/v) containing 10 mM ammonium formate. The gradient commenced at 85% A/15% B. Mobile phase B was increased to 50% at 2.5 min, 57% at 2.6 min, 70% at 9.0 min, 93% at 9.1 min, 96% at 11.0 min and 100% at 11.1 min. The column was maintained at 100% B until 12.0 min, returned to 85% A/15% B at 12.2 min, and maintained under the initial conditions until the end of the 16-min run.

Mass spectrometric data were acquired in positive- and negative-ion modes using dynamic multiple-reaction monitoring (dMRM). The core MRM methodology and lipid-specific transitions were based on our previously described targeted lipidomics platform^86^, with additional transitions incorporated from the expanded high-sensitivity cellular lipidomics panel^87^ to provide broader lipid-species coverage.

Source conditions were: drying-gas temperature, 200 °C; drying-gas flow, 17 L/min; nebuliser pressure, 20 psi; sheath-gas temperature, 280 °C; and sheath-gas flow, 10 L/min. Capillary voltages were 3500 V and 3000 V in positive- and negative-ion modes, respectively, and nozzle voltages were 1000 V and 1500 V, respectively. Positive-ion high- and low-pressure iFunnel RF voltages were 200 and 110 V, respectively, while the corresponding negative-ion values were 150 and 60 V. The LC effluent was diverted to waste outside the scheduled dMRM acquisition period and from 13.1 min onward.

#### Phosphoinositide analysis

Targeted LC-MS/MS analysis of phosphoinositides was performed using an Agilent 1290 Infinity II LC system coupled to an Agilent 6495C triple-quadrupole mass spectrometer equipped with an Agilent Jet Stream electrospray ionisation source. Chromatographic separation was performed using a Waters ACQUITY BEH C8 Vanguard Premier column (2.1 x 100 mm) maintained at 55 °C. Samples (5 µL) were injected at a flow rate of 0.30 mL/min, with the autosampler needle washed for 10 s at the flush port following each injection.

Mobile phase A consisted of 50:50 acetonitrile/water (v/v) containing 20 mM ammonium bicarbonate and 5 µM medronic acid, while mobile phase B consisted of 80:10:10 isopropanol/acetonitrile/water (v/v/v) containing 10 mM ammonium bicarbonate and 2.5 µM medronic acid. The gradient commenced at 100% A and was changed to 90% A/10% B at 1.0 min, 55% A/45% B at 3.0 min and 25% A/75% B at 15.0 min. Mobile phase B was increased to 100% at 16.0 min and maintained until 17.5 min. The system was returned to 100% A at 18.0 min and maintained under these conditions until the end of the 20-min run. A targeted list of phosphoinositides was acquired in positive ion mode corresponding to the [M+H]^+^ and [M+NH4]^+^ ion of each species.

Mass spectrometric acquisition was performed in positive-ion mode using dMRM. Source conditions were: drying-gas temperature, 200°C; drying-gas flow, 15 L/min; nebuliser pressure, 20 psi; sheath-gas temperature, 320°C; sheath-gas flow, 12 L/min; capillary voltage, 4500 V; and nozzle voltage, 1500 V. Positive-ion iFunnel RF voltages were 210 V for the high-pressure funnel and 110 V for the low-pressure funnel.

## Data availability

Atomic coordinates were deposited in the Protein Data Bank (PDB) under accession codes 47OL (FZD5 in complex with Gq in LMNG micelle), 47OM (FZD7 in complex with Gs in LMNG micelle), 47ON (FZD7 in complex with Gs in lipid nanodisc), 47OO (FZD7 in complex with Gs in GDN micelle) and 47OP (Quad Mutant FZD7 in complex with Gs in LMNG micelle). Cryo-EM maps were deposited in the Electron Microscopy Data Bank under the accession codes EMD-84688 (Non-Uniform refinement map of human Frizzled 5 in complex with mini Gs in LMNG micelle), EMD-84689 (Composite map of human Frizzled 7 in complex with mini Gs in LMNG micelle), EMD-84492 (Non-uniform refinement map of human Frizzled 7 in complex with mini Gs in LMNG micelle), EMD-84493 (Local refinement map of human Frizzled 7 in complex with mini Gs in LMNG micelle), EMD-84690 (Composite map of human Frizzled 7 in complex with mini Gs in lipid nanodisc), EMD-84489 (Non-uniform refinement map of human Frizzled 7 in complex with mini Gs in lipid nanodisc), EMD-84490 (Local refinement map of human Frizzled 7 in complex with mini Gs in lipid nanodisc), EMD-84691 (Non-Uniform refinement map of human Frizzled 7 in complex with mini Gs in GDN micelle), EMD-84692 (Composite map of mutant human Frizzled 7 in complex with mini Gs in LMNG Micelle), EMD-84494 (Non-Uniform Refinement map of mutant human Frizzled 7 in complex with mini Gs in LMNG Micelle), EMD-84495 (Local Refinement map of mutant human Frizzled 7 in complex with mini Gs in LMNG Micelle).

## Acknowledgements

This work was supported by the Monash University Ramaciotti Centre for cryo-electron microscopy, the Ian Holmes Imaging Centre, Bio21 Institute, the Monash University MASSIVE high-performance computing facility and supercomputing resources, and the National Computational Merit Allocation Scheme (NCMAS), with access to computational resources provided by the National Computational Infrastructure (NCI). ΔDVL1-3 HEK293T cells were a kind gift from Stephane Angers (University of Toronto). ΔFZD1-10 HEK293T cells were a kind gift from Benoit Vanhollebeke (Université libre de Bruxelles). nLuc tagged β_2_AR was a kind gift from Carl White (University of Melbourne). TRUPATH constructs were generously provided by Dr. Bryan Roth (University of North Carolina). Venus-WT Gα_s_, Gα_s_, Gβ_1_, and Gγ_2_–Venus were a kind gift from Peishen Zhao (Monash University). The authors thank Hannes Schihada (Marburg University) for validating WT FZD5 and WT FZD7 constructs. The authors acknowledge the WEHI Protein Production Facility for purification of the FZD7-mGs complex in GDN detergent micelle. The AI tool Claude Sonnet 4.6 (Anthropic, 2026) was used for proofreading the manuscript and checking for consistency in spelling, grammar, and clarity of the text. It was not used for the generation of ideas, content, figures, or data.

## Author contributions

W.A.C.B. and A.G. designed the overall research. W.A.C.B., S.D., J.R.P. and A.G. designed, expressed, and purified protein samples. W.A.C.B., S.D., H.V., N.S.K., A.P.L., A.G. performed sample vitrification and cryo-EM imaging. W.A.C.B., S.D. and A.G. processed the cryo-EM data and generated and analysed atomic models. W.A.C.B., A.S.Q., B.J.W-N. and M.L.O. designed, performed, and analysed MD simulations. W.A.C.B., S.D., L.L.L.W., T.T.N. generated DNA constructs and performed and analysed pharmacology experiments. G.S. provided and validated DNA constructs. T.J. and K.H. performed and analysed lipidomics experiments. W.A.C.B., M.E.C., G.S., M.L.O. and A.G. provided data review and supervision. W.A.C.B. and A.G. wrote the manuscript with contributions and discussions from all authors.

