## Supplementary Information for "Structural basis for inefficient G protein coupling by Frizzled receptors"

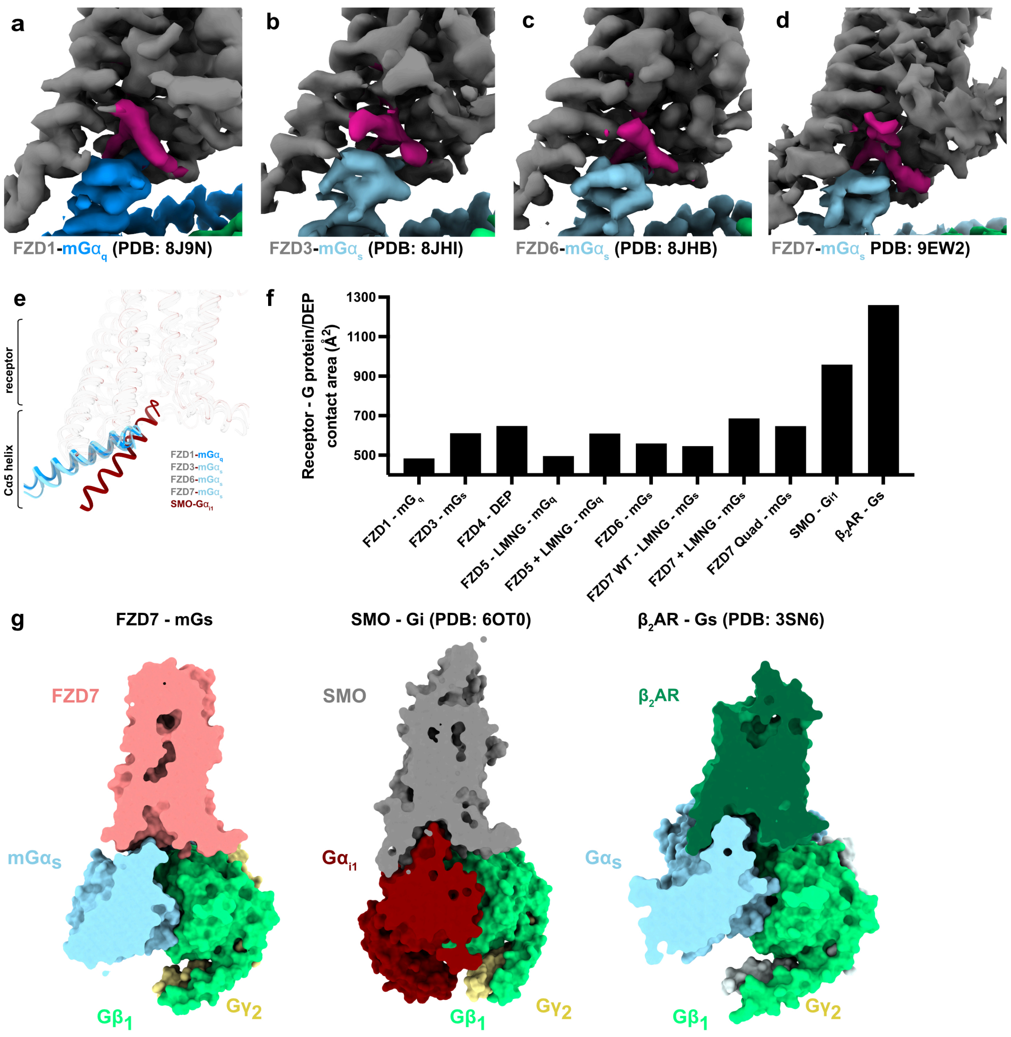


**Supplementary Fig. 1 | Lipid- or detergent-like density at the intracellular receptor-G protein interface is a recurring feature in FZD-G protein structures.** Close-ups of the intracellular receptor-G protein interface in published cryo-EM maps, showing unassigned density (magenta) between the receptor (grey) and the Gα subunit: **a** FZD1-mGα$_q$ (PDB 8J9N, contour level 0.118, σ = 4.80). **b** FZD3-mGα$_s$ (PDB 8JHI, contour level 0.118, σ = 6.07). **c** FZD6-mGα$_s$ (PDB 8JHB, contour level 0.067, σ = 2.56). **d** FZD7-mGα$_s$ (PDB 9EW2, contour level 0.016, σ = 4.55). **e** Superposition of the Gα α5 helices from the FZD1-mGα$_q$, FZD3-mGα$_s$, FZD6-mGα$_s$ and FZD7-mGα$_s$ complexes with SMO-Gα$_{i1}$, aligned on the receptor transmembrane domain. **f** Buried surface area between the receptor and the G protein or DEP domain (Å$^2$), calculated from the indicated structures with PISA. Each bar is a single value calculated from one structure; no error is associated with these values. **g** Cross-sections through the molecular surfaces of FZD7-mGs, SMO-Gi1 (PDB 6OT0) and β$_2$AR-Gs (PDB 3SN6), illustrating the difference in G protein engagement with the receptor TM core.

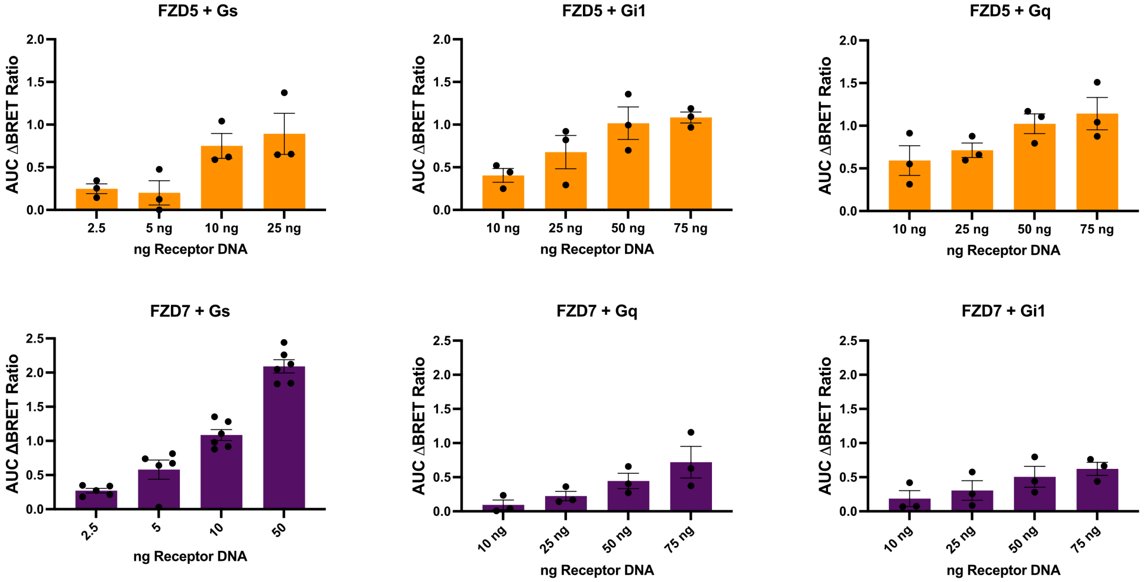


**Supplementary Fig. 2 | FZD5 and FZD7 both activate Gs, Gi1 and Gq heterotrimer in a receptor-expression-dependent manner.** ΔFZD1-10 HEK293T cells were transfected with TRUPATH BRET biosensors and increasing amounts of WT FZD5 or WT FZD7. Constitutive G protein activation measured by TRUPATH BRET, expressed as AUC of the ΔBRET ratio. Bars show mean ± s.e.m. with individual points representing independent experiments performed in triplicate. The FZD7-mGs data are the constitutive activity data of Fig. 4d replotted as bar graphs. Source data are provided as a Source Data file.

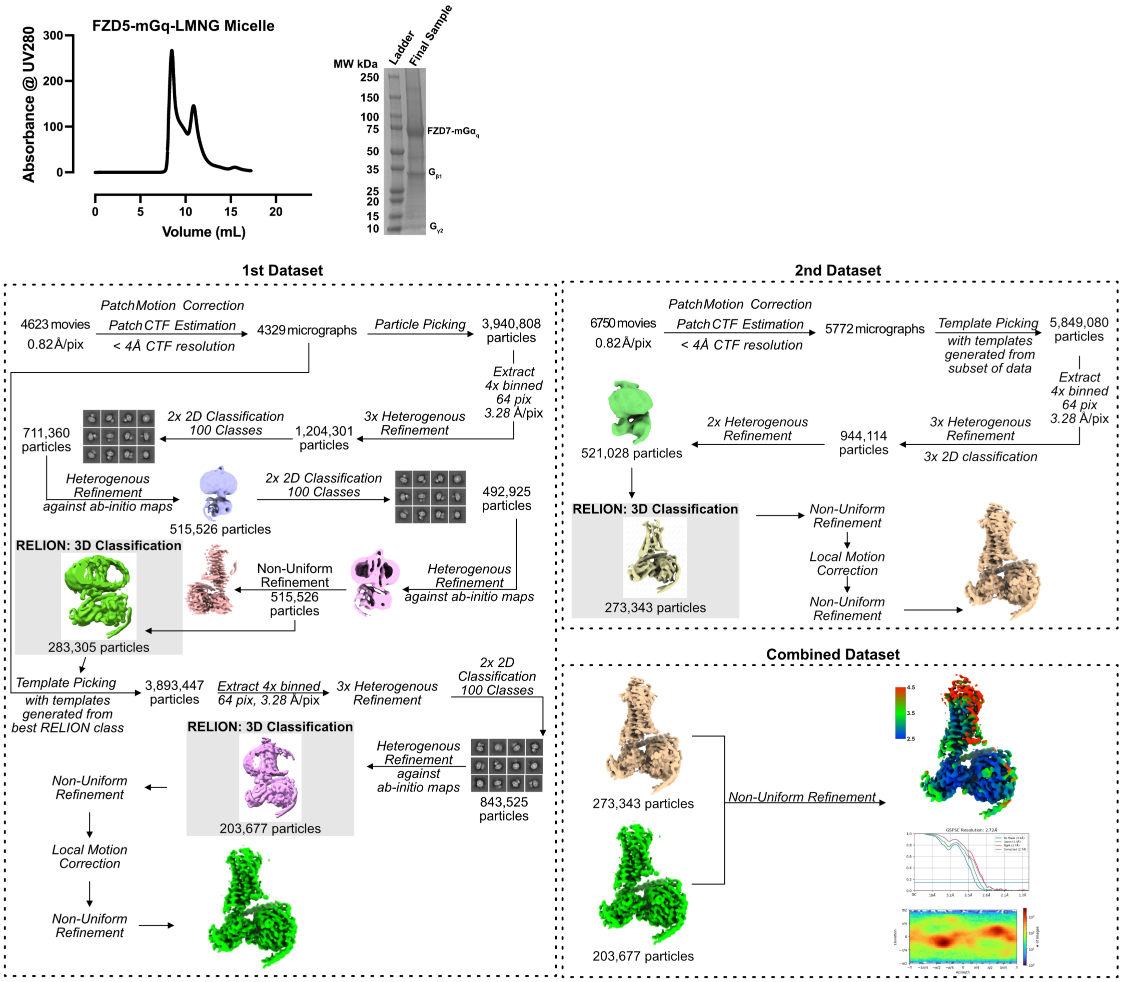


**Supplementary Fig. 3 | Purification and cryo-EM data processing for the FZD5-mGq complex in LMNG micelle.** Flow chart for cryo-EM analysis as described in methods leading to final cryo-EM density maps for FZD5-mGq complex in LMNG micelle.

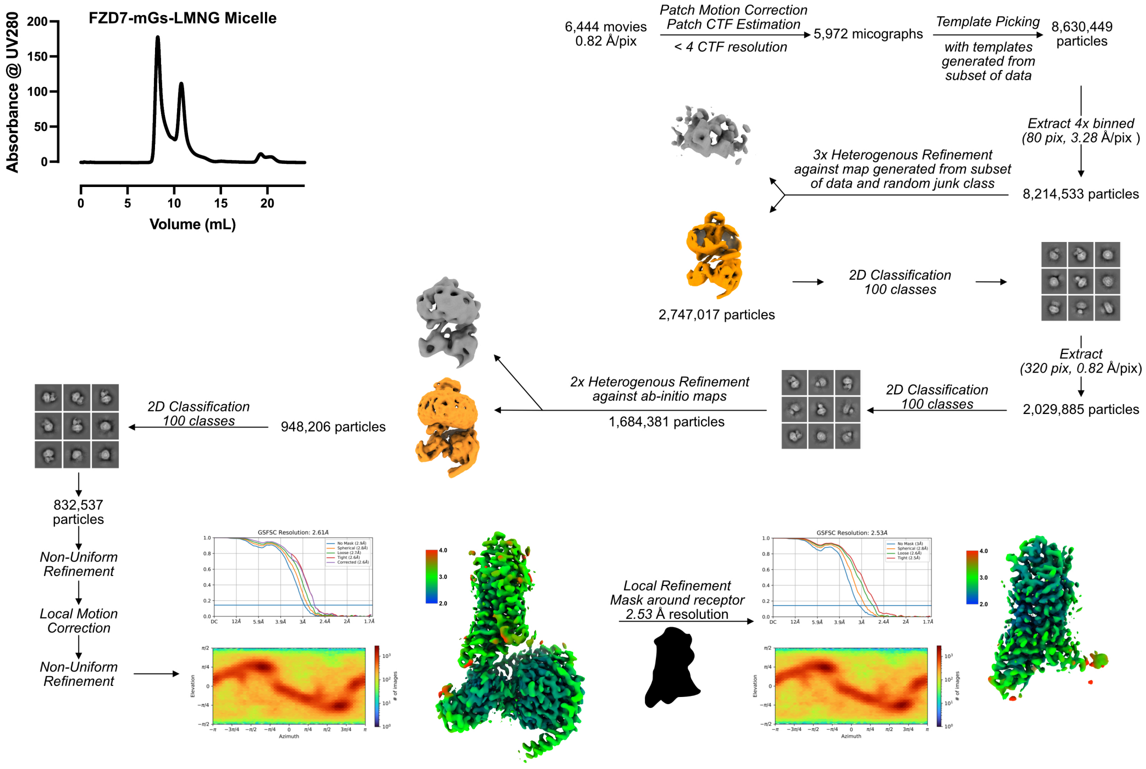


**Supplementary Fig. 4 | Purification and cryo-EM data processing for the FZD7-mGs complex in LMNG micelle.** Flow chart for cryo-EM analysis as described in methods leading to final cryo-EM density maps for FZD7-mGs complex in LMNG micelle.

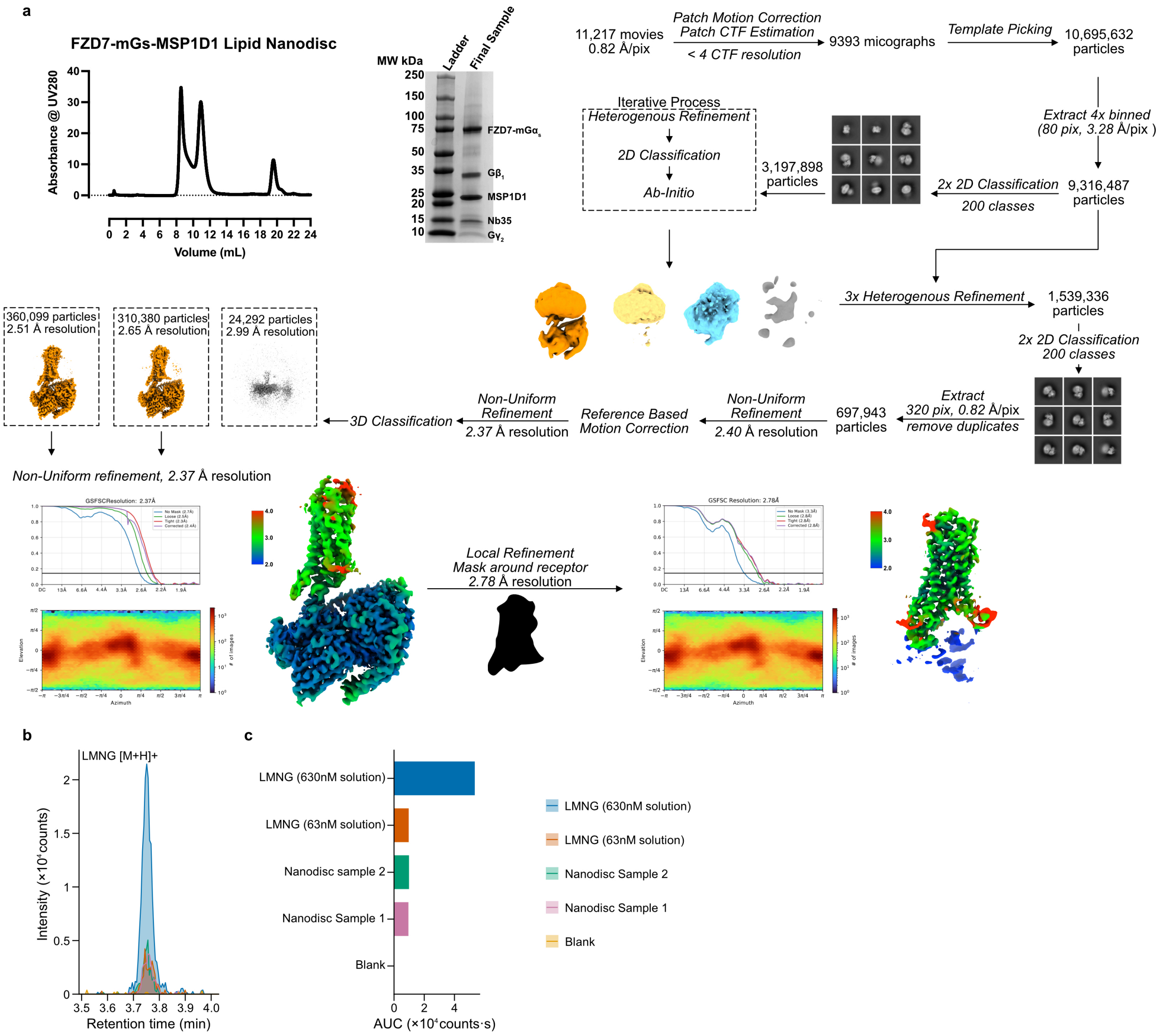


**Supplementary Fig. 5 | Purification and cryo-EM data processing for the FZD7-mGs complex in MSP1D1 lipid nanodisc. a** Flow chart for cryo-EM analysis as described in methods leading to final cryo-EM density maps for FZD7-mGs complex in MSP1D1 lipid nanodisc. **b** Representative liquid chromatography–mass spectrometry (LC–MS) extracted-ion chromatograms of protonated LMNG ($[M+H]^+$) in two FZD7–mGs nanodisc samples (Nanodisc 1 and 2), blank, and LMNG reference solutions (63 and 630 nM). LMNG-associated signals were detected at approximately 3.7 min in both nanodisc samples, with retention times corresponding to the reference solutions. **c** Integrated chromatographic peak areas (AUC; counts·s) for the LMNG signal in each sample and reference solutions.

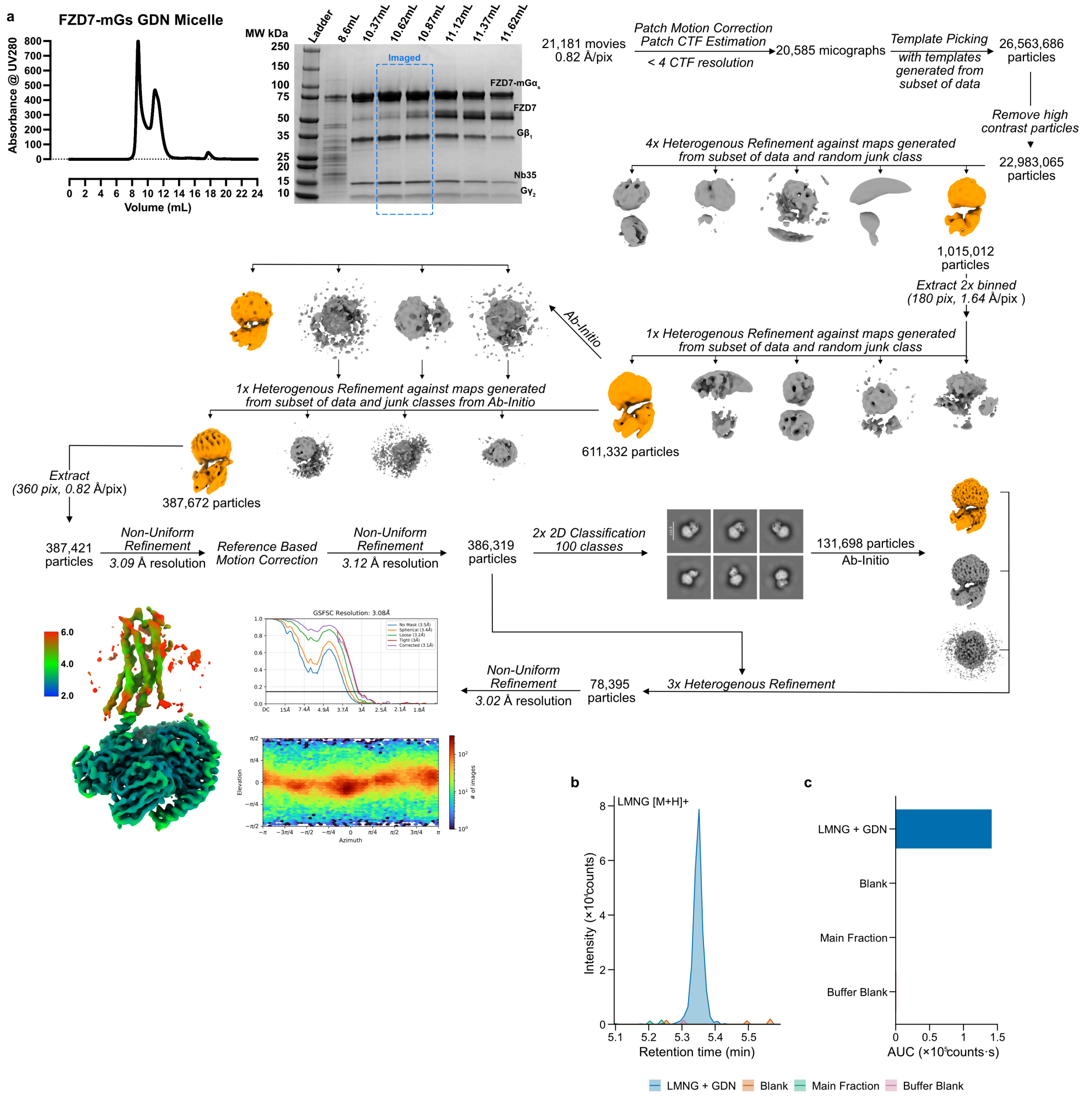


**Supplementary Fig. 6 | Purification and cryo-EM data processing for the FZD7-mGs complex in GDN micelle. a** Flow chart for cryo-EM analysis as described in methods leading to final cryo-EM density maps for FZD7-mGs complex in GDN micelle. **b** Representative LC–MS extracted-ion chromatograms monitoring protonated LMNG ($[M+H]^+$) in the main FZD7–mGs protein fraction following purification using GDN, alongside a purification-buffer blank, a separate blank and an LMNG-spiked GDN reference. The LMNG-spiked GDN reference produced a distinct chromatographic peak at approximately 5.3 min, whereas no corresponding signal above background was detected in the purified protein fraction or blank samples. **c** Integrated LMNG chromatographic peak areas for each sample.

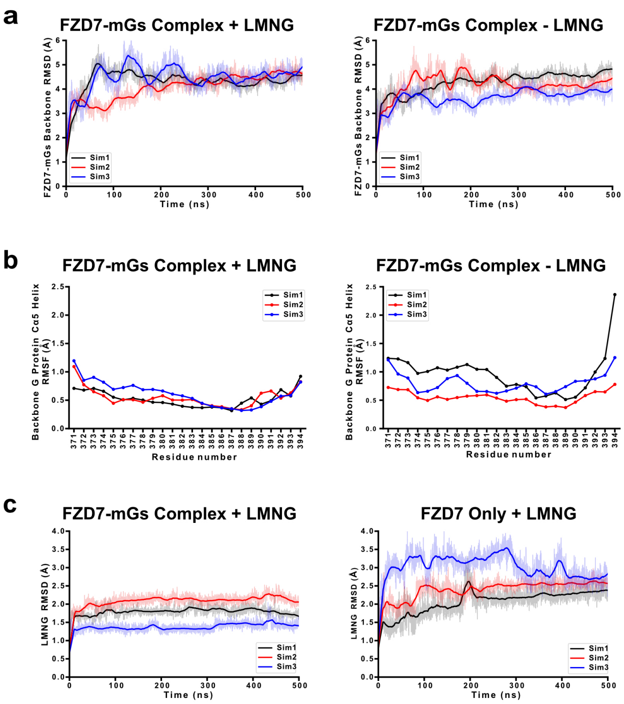


**Supplementary Fig. 7 | Stability of the FZD7-mGs complex and of bound LMNG during molecular dynamics simulations. a** Backbone root mean square deviation (RMSD) of the FZD7-mGs complex over 500 ns of simulation with LMNG bound (left) and after removal of LMNG (right). **b** Backbone RMSF of the $G\alpha_s$ α5 helix (residues 371-394) for the same simulations. **c** RMSD of the LMNG molecule in the FZD7-mGs complex (left) and in simulations of FZD7 receptor alone (right). In all panels, three independent trajectories per condition are shown in different colours.

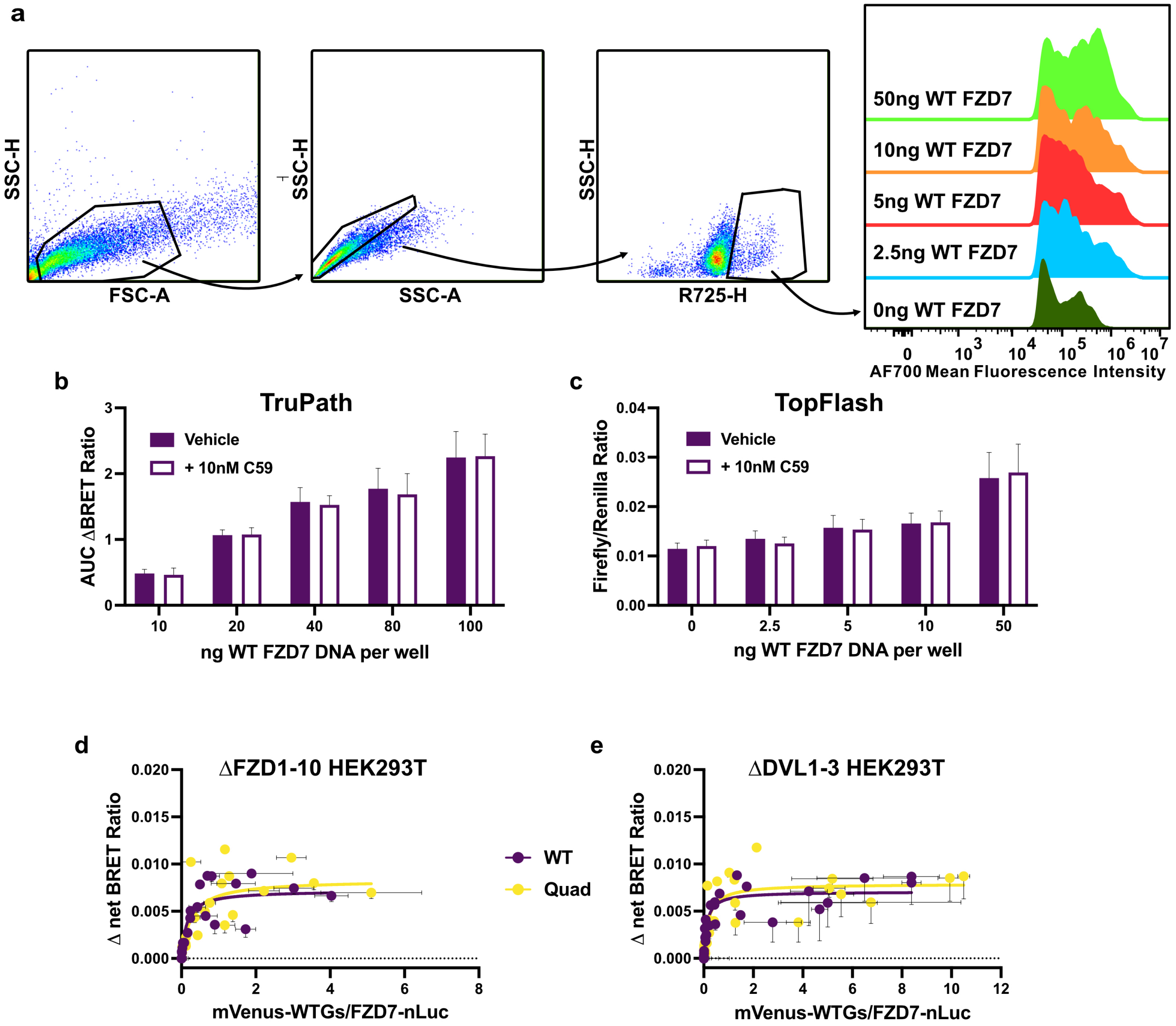


**Supplementary Fig. 8 | Receptor expression, Wnt-independence of FZD7 constitutive activity, and DVL-independent Gs coupling.** **a** Representative flow cytometry gating strategy used to quantify cell-surface receptor expression: successive gates on HEK293T cells (FSC-A/SSC-H), single cells (SSC-A/SSC-H) and FLAG +ve cells (R725-H/SSC-H), followed by AF700 mean fluorescence intensity histograms for ΔFZD1-10 HEK293T cells transfected with 0, 2.5, 5, 10 and 50 ng WT FZD7 DNA. **b** Constitutive Gs activation (AUC of the ΔBRET ratio) in ΔFZD1-10 HEK293T cells transfected with the indicated amounts of WT FZD7 DNA and treated with vehicle (filled bars) or 10 nM of the Porcupine inhibitor C59 (open bars). **c** TOPFlash reporter activity (firefly/Renilla ratio) under the same conditions as b. In b and c, bars show mean ± s.e.m. from n = 3 independent experiments. **d-e** BRET saturation assay of Gs recruitment in ΔFZD1-10 HEK293T cells (**d**) and ΔDVL1-3 HEK293T (**e**) cells, transfected with constant receptor-nLuc (donor) and increasing mVenus-WT $G\alpha_s$ (acceptor); net BRET ratio plotted against mVenus Fluorescence/nLuc luminescence ratio. Data points represent mean ± s.e.m. (both axes), pooled from n = 7 (ΔFZD1-10) and n = 8 (ΔDVL1-3) independent experiments performed in duplicate for WT FZD7 and Quad. Curves represent one-site specific binding fits. Parameters are listed in Supplementary Table 3. Source data are provided as a Source Data file.

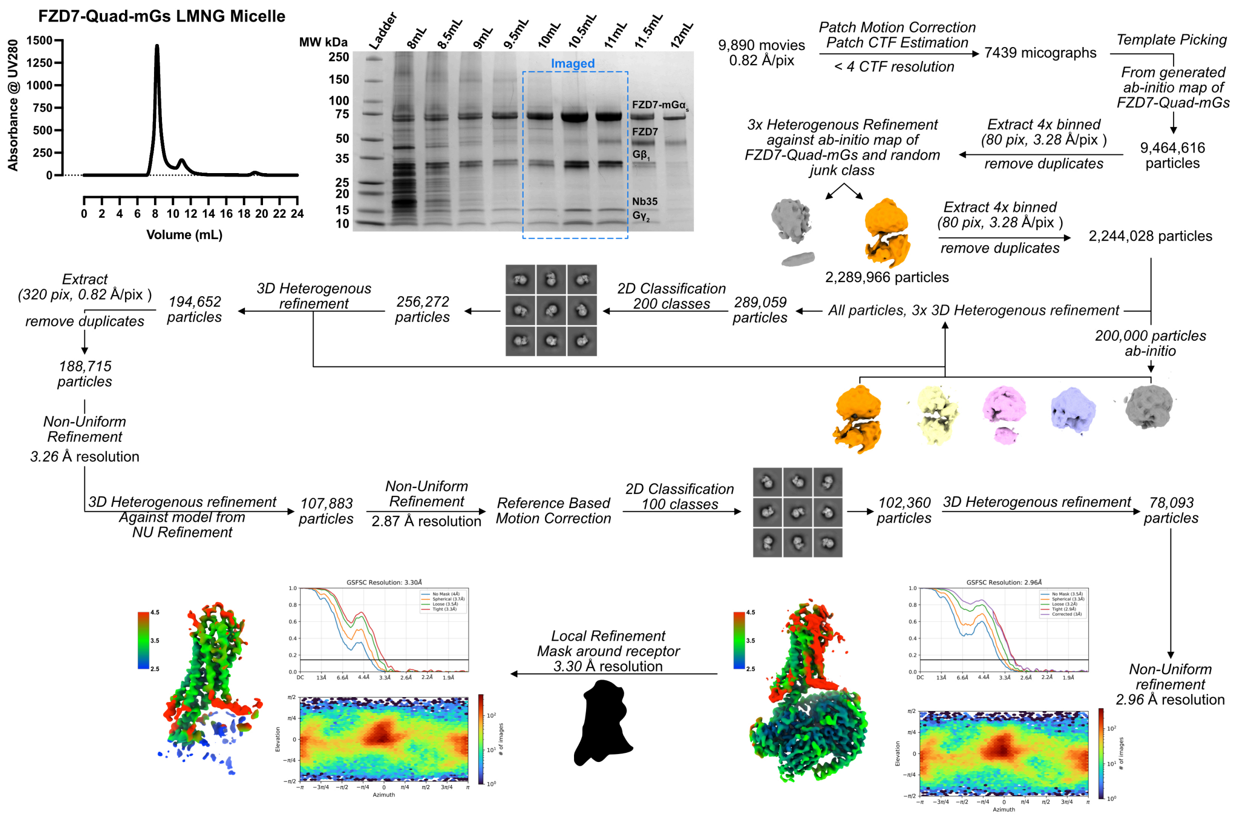


**Supplementary Fig. 9 | Purification and cryo-EM data processing for the FZD7-Quad-mGs complex in LMNG micelle.** Flow chart for cryo-EM analysis as described in methods leading to final cryo-EM density maps for FZD7 Quad mutant-mGs complex in LMNG micelle.

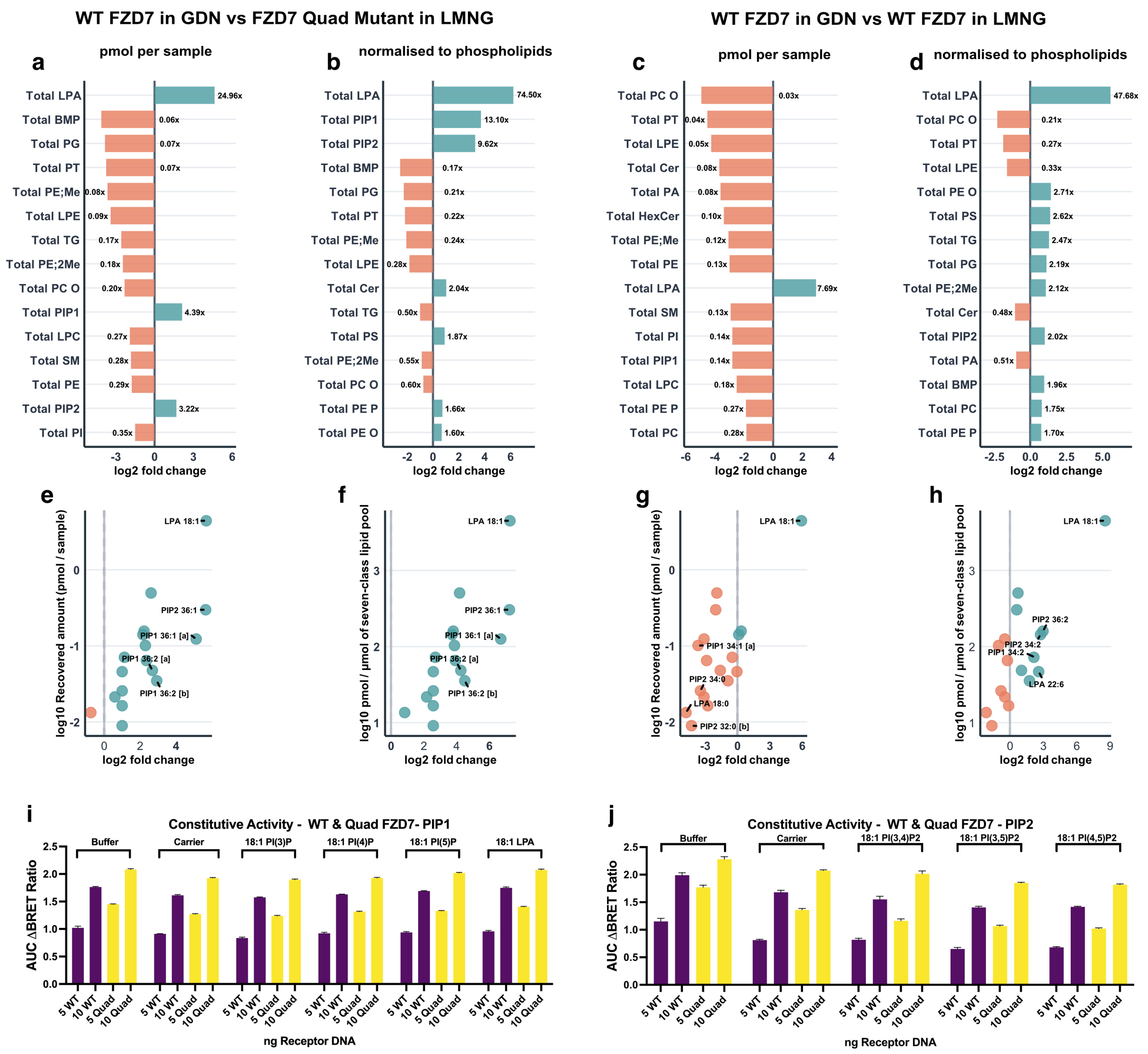


**Supplementary Fig. 10 | Lipid enrichment in purified FZD7 complexes and the effect of exogenous phosphoinositides on FZD7 constitutive activity.** Lipidomics was performed on WT FZD7–mGs complexes purified in GDN or LMNG and the FZD7 Quad mutant (E286$^{2.41}$A/A365$^{3.54}$F/K466$^{6.28}$E/L467$^{6.29}$T) purified in LMNG to investigate potential endogenous lipids associated with the receptor–G protein complex. **a-d** Relative differences in recovered lipid class abundances between WT FZD7–mGs purified in GDN and either the FZD7 Quad mutant purified in LMNG (a, b) or WT FZD7–mGs purified in LMNG (c, d). Lipid class abundances were compared as absolute recovered amounts (pmol per sample; a, c) or following normalisation to the combined abundance of seven reference lipid classes (PC, PA, SM, PE, PG, PI and PS; pmol per µmol of reference lipid pool; b, d). The 15 lipid classes exhibiting the largest absolute $\log_2$ fold changes after blank-based quality screening are displayed. Bars represent $\log_2$ fold changes calculated as GDN-purified WT FZD7 relative to the corresponding LMNG-purified comparator, with teal indicating higher and coral indicating lower abundance in the GDN preparation. Fold-change values are annotated alongside each bar. e-h Lipid species-level comparisons focusing on lysophosphatidic acids (LPA), monophosphorylated phosphatidylinositols (PIP1) and bisphosphorylated phosphatidylinositols

(PIP2). Each point represents an individual annotated lipid species passing the matched GDN-blank signal screen, plotted according to its $\log_2$ fold change between purification conditions (x-axis) and $\log_{10}$ abundance in the GDN-purified WT FZD7 sample (y-axis). Abundances are expressed as recovered pmol per sample (e, g) or pmol per µmol of the seven-class reference lipid pool (f, h). Comparisons are shown against the LMNG-purified Quad mutant (e, f) and LMNG-purified WT FZD7 (g, h). Selected lipid species, including LPA 18:1 and individual PIP1/PIP2 species, are annotated. Lipid quantities were derived from background-corrected measurements, with candidate lipid species screened against a matched GDN blank. Each condition represents one analysed pooled protein preparation over multiple LC-MS runs. **i, j** ΔFZD1-10 HEK293T cells were transfected with TRUPATH BRET biosensors and 0, 5 or 10 ng per well of WT FZD7 DNA. After 48h, cells were treated with lipids complexed with lipid carrier (described in methods). Constitutive G protein activation measured by TRUPATH BRET, expressed as AUC of the ΔBRET ratio. Bars show mean ± s.d. of one individual independent experiment performed in duplicate. Source data are provided as a Source Data file.

**Supplementary Table 1 | Data collection and refinement statistics**

| **Data Collection** | **FZD5-mG$_{sqNi}$/Nb35 (LMNG Micelle)** | **FZD7-mG$_s$/Nb35 (LMNG Micelle)** | **FZD7-mG$_s$/Nb35 (Lipid Nanodisc)** | **FZD7-mG$_s$/Nb35 (GDN Micelle)** | **FZD7 Quad Mutant - mG$_s$/Nb35 (LMNG Micelle)** |
|---|---|---|---|---|---|
| **Micrographs** | 11,373 | 6,444 | 11,217 | 21,181 | 9,890 |
| **Electron dose ($e^-$/Å$^2$)** | 60 | 60 | 60 | 60 | 60 |
| **Voltage (kV)** | 300 | 300 | 300 | 300 | 300 |
| **Pixel size (Å)** | 0.82 | 0.82 | 0.82 | 0.82 | 0.82 |
| **Defocus range (μM)** | 0.5 - 1.2 | 0.5 - 1.2 | 0.5 - 1.2 | 0.5 - 1.2 | 0.5 - 1.2 |
| **Symmetry Imposed** | C1 | C1 | C1 | C1 | C1 |
| **Particles (final map)** | 477,020 | 832,537 | 670,479 | 78,395 | 78,093 |
| **Resolution (0.143 FSC) (Å)** | 2.72 | 2.61 | 2.78 | 3.02 | 2.96 |
| **Refinement** | | | | | |
| **CCmap_model** | 0.80 | 0.85 | 0.87 | 0.72 | 0.79 |
| **Model Quality** | | | | | |
| **R.m.s deviations** | | | | | |
| **Bond Length (Å)** | 0.003 | 0.004 | 0.004 | 0.004 | 0.004 |
| **Bond angles (°)** | 0.649 | 0.639 | 0.620 | 0.611 | 0.639 |
| **Ramachandran** | | | | | |
| **Favoured (%)** | 98.62 | 98.75 | 98.43 | 98.43 | 98.53 |
| **Outliers (%)** | 0 | 0 | 0 | 0 | 0 |
| **Rotamer outliers (%)** | 0.49 | 0.84 | 0.75 | 0.75 | 0.85 |
| **C-Beta deviations (%)** | NA | NA | NA | NA | NA |
| **Clashscore** | 5.21 | 5.32 | 4.57 | 15.69 | 4.44 |
| **MolProbity score** | 1.28 | 1.29 | 1.23 | 1.70 | 1.22 |

**Supplementary Table 2 | Functional parameters for Wnt-3A in Topflash Assays**

| | WT FZD7 | E286$^{2.41}$A | A365$^{3.54}$F | K466$^{6.28}$E | L467$^{6.29}$T | Quad Mutant |
|---|---|---|---|---|---|---|
| **$EC_{50}$ (ng/mL)[a]** | 993.9 ± 131.17 (4) | 851.5 ± 71.16 (4) | N.R. | 1002.3 ± 219.21 (3) | N.R. | N.R. |
| **$E_{max}$[b]** | 114.2 ± 4.08 (4) | 89.3 ± 21.92 (4) | N.R. | 46.6 ± 14.41 (3)* | N.R. | N.R. |

Data represent the mean ± S.E.M. of (n) individual experiments performed in duplicate. * significantly different from WT, one-way ANOVA, Dunnett's post-hoc test, ($P < 0.05$). N.R. no response.

[a] Concentration of Wnt-3A (ng/mL) required to give half maximal response obtained from four-parameter logistic equation.

[b] Maximal response for Wnt-3A obtained from four-parameter logistic equation.

**Supplementary Table 3 | Functional parameters for net BRET saturation binding assays**

| | WT FZD7 | Quad Mutant | $\beta_2$AR |
|---|---|---|---|
| | *Mini GαS in ΔFZD$_{1-10}$ HEK293T Cells* | | |
| **$BRET_{50}$**[a] | 1.25 ± 0.27 (4) | 1.69 ± 0.26 (3) | 1.52 ± 0.28 (3) |
| **$BRET_{Max}$**[b] | 0.018 ± 0.0018 (4) | 0.020 ± 0.0015 (3) | 0.040 ± 0.0023 (3)* |
| | *mVenus-DVL2 in ΔFZD$_{1-10}$ HEK293T Cells* | | |
| **$BRET_{50}$**[a] | 0.59 ± 0.11 (4) | N.R. | N.R. |
| **$BRET_{Max}$**[b] | 0.019 ± 0.0019 (4) | N.R. | N.R. |
| | *WT GαS in ΔFZD$_{1-10}$ HEK293T Cells* | | |
| **$BRET_{50}$**[a] | 0.14 ± 0.050 (7) | 0.26 ± 0.094 (7) | N.D. |
| **$BRET_{Max}$**[b] | 0.0072 ± 0.0005 (7) | 0.0083 ± 0.0007 (7) | N.D. |
| | *WT GαS in ΔDVL$_{1-3}$ HEK293T Cells* | | |
| **$BRET_{50}$**[a] | 0.12 ± 0.037 (9) | 0.18 ± 0.054 (9) | N.D. |
| **$BRET_{Max}$**[b] | 0.0071 ± 0.0004 (9) | 0.0079 ± 0.0005 (9) | N.D. |

Data represent the mean ± S.E.M. of (n) pooled, independent experiments performed in duplicate. N.R. no response. N.D. not determined. * Significantly different to WT FZD7. Differences in $BRET_{50}$ and $BRET_{max}$ between WT FZD7 and Quad Mutant or WT FZD7 and $\beta_2$AR were assessed using an extra sum-of-squares F test. *$p < 0.05$.

[a] $BRET_{50}$ is the acceptor/donor ratio at which half $BRET_{Max}$ is reached and reflects relative affinity as obtained from a one-site specific binding model.

[b] $BRET_{Max}$ is the maximal net BRET ratio as obtained from a one-site specific binding model.
